# Multi-protein assemblies orchestrate co-translational enzymatic processing on the human ribosome

**DOI:** 10.1101/2024.06.14.599006

**Authors:** Marius Klein, Klemens Wild, Irmgard Sinning

## Abstract

Most nascent chains undergo rapid co-translational enzymatic processing as soon as their N-terminus becomes accessible at the ribosomal polypeptide tunnel exit (PTE). In eukaryotes, N-terminal methionine excision (NME) by Methionine Aminopeptidases (MAP1 and MAP2), and N-terminal acetylation (NTA) by N-Acetyl-Transferase A (NatA), is the most common set of subsequent modifications carried out on the ribosome. How these two enzymatic processes are coordinated in quick succession in the context of a rapidly translating ribosome has remained elusive.

Here, we report that human NatA occupies a non-intrusive ‘distal’ binding site on the ribosome which does not interfere with most other ribosome associated factors (RAFs). In this position, NatA can partake in a coordinated and dynamic assembly with MAP1 through a complex scaffolding function of the abundant Nascent Polypeptide Associated Complex (NAC). Alternatively, MAP2 can co-occupy the PTE with NatA in preparation for successive NME and NTA. In contrast to MAP1, MAP2 completely covers the PTE and is thus incompatible with NAC and MAP1 recruitment. Both assemblies can compile on the human ribosome independent of nascent chain substrates. Together, our structures provide the structural framework for the rapid coordinated orchestration of NME and NTA in protein biogenesis.

## Introduction

At early states of protein biogenesis, the N-terminus of the nascent chain experiences vivid competition between different ribosome associated factors (RAFs) which stand in line to cleave, acetylate or myristoylate these residues^1,2^. When the N-terminus starts to protrude from the polypeptide tunnel exit (PTE), the initiator methionine is in a highly predictable position while the nascent chain remains confined within the exit tunnel. This unique opportunity is most prominently exploited by methionine aminopeptidases and acetylases, which must carry out N-terminal methionine excision (NME) and N-terminal acetylation (NTA) in a coordinated manner. In eukaryotes the vast majority of proteins experiences co-translational NME, followed by NTA^1^. Both modifications are crucial, as they impinge on protein turnover, function, localization and stability^3^. Misfunction in either machinery has been implicated in disease^4–8^.

Strikingly, Methionine Aminopeptidases (MAPs) and N-terminal Acetyltransferases (NATs) are low in abundance relative to the number of ribosomes inside the cell^9^. Their underrepresentation necessitates well-coordinated regulation over their timely recruitment, function and eventual displacement. The high demand for co-translational NME is attended to by two catalytically similar enzymes in eukaryotes. The two eukaryotic MAP isoforms, MAP1 and MAP2 both share the characteristic pita-bread fold with their active dinuclear centres^10^. However, specific sequence insertions in MAP2 (including a prominent ∼60 AS insert domain), as well as fundamental differences in the architecture of the N-terminal extension, elicit changes in how these factors are recruited to and accommodated on the ribosomal surface. The zinc-finger motive containing N-terminus of mammalian MAP1 is recognized by the nascent polypeptide associated complex (NAC), which administers MAP1 recruitment to the PTE^11^. In contrast, our recent structures of MAP2 reveal a different, factor independent mode of binding to the ribosome, and a more central placement on the PTE, as well as the presence of a second binding site in the ribosomal A-site^12^. Beyond its role as a co-translational protease, MAP2 has been shown to influence translational initiation^13^ and might also act on other stages of the translational cycle via its A-site interaction^12^.

The modular co-translational machinery dedicated to the N-terminal acetylation (NTA) of nascent chain substrates encompasses five conserved N-Acetyltransferases (NATs) (NatA-E) with distinct substrate specificities^3^. The heterodimeric NatA complex carries out the majority of acetylation events and possesses the broadest substrate selectivity (Gly-, Ala-, Ser-, Thr-, Val-, Cys-). Unlike NatB or NatC, NatA cannot acetylate the initiator methionine, thus necessitating prior NME activity by MAP1 or MAP2^3^. As these processes must coincide within a narrow window of opportunity while translation is rapidly progressing, a functional synchronisation between NME and NTA appears likely. The ribosome structure with the NatA containing heterotrimer NatE has been determined in yeast^14^. Given that the third subunit Naa50 strongly contributes to the 80S interaction, the absence of this subunit might affect how NatA engages with the ribosome.

In addition to Naa50, HypK has been identified as an important NatA interactor, which binds to the α-helical Naa15 scaffold^15^. HypK composes of a three-helix bundle UBA like domain, a long central helix and a mostly disordered extended N-terminus. In complex with NatA, HypK has a strongly stabilizing effect and inhibits the catalytic activity of Naa10^16,17^. Moreover, HypK binding to NatA induces structural rearrangements in Naa15, which allosterically inhibits Naa50 binding^18^. HypK has been reported to co-localize with polysomes^15^ and contact nascent chains^19^ while loss of HypK function negatively impacts NatA function^15,20,21^. While the mechanisms of *in vitro* NatA inhibition by HypK are well characterized^16–18^, the molecular details of NatA activation *in vivo* remain vague. The C-terminus of HypK, including the UBA domain, is homologous to the C-terminus of NACα. However, the precise functional link between these proteins remains to be elucidated.

Aside from ribosomal proteins and the ubiquitous rRNA, the mostly unstructured heterodimer NAC also shapes and influences the biochemical properties of the PTE. Its large surface provided by the four charged and extended protein termini might have far-reaching implications on other RAFs that associate to the tunnel exit during translation. Work on NAC has already highlighted its importance in SRP regulation^22–25^, protein folding and proteostasis^26^ as well as MAP1 coordination^11,27^. Its omnipresence in this early stage of co-translational protein maturation strongly implies an additional role in the regulation of acetylation.

In this work we report the cryo-EM structure of NatA on the human 80S ribosome. Unlike yeast NatE, human NatA binds at a non-intrusive ‘distal’ position which could allow concurrent binding of MAP1, MAP2, Ebp1, NAC, NatB, RAC, SRP or SEC61 without steric clash. In its distal position, NatA can bind together with MAP2 to form a ternary complex with the ribosome, capable of successive NME and NTA. Alternatively, NatA can form a dynamic ring-like assembly with MAP1 and NAC which encompasses the PTE. Overall, our structures provide evidence that RAFs can arrange in dynamic multi-protein PTE assemblies independent of nascent chains, in preparation for co-translational enzymatic processing on the human ribosome.

## Results

### Human NatA occupies a ‘distal’ position on the ribosome

The heterodimeric NatA complex comprises of the large scaffold protein Naa15, and the small Naa10 enzyme with the characteristic GNAT (GCN5-related-N-acetyltransferase) fold, characterized by a conserved α/μ-fold^28–30^. Herein, Naa15 composes of 13 tetratricopeptide repeat (TPR) motifs (45 α-helices) which form an elaborate ring-like scaffold that engulfs the acetyltransferase^16^. In *Saccharomyces cerevisiae (Sc)*, the NatA complex is exclusively present as a heterotrimer with a catalytically inactive Naa50^31^. Given that Naa50 strongly contributes to ribosome binding of *Sc*NatA^14^, its absence in human NatA should affect this interaction. One major function of Naa15 is to facilitate the positioning of Naa10 near the PTE during translation^14,32–34^. To examine how this is accomplished at the human ribosome, we reconstituted the NatA-80S complex *in vitro* and subjected the sample to single particle (SPA) cryo-EM analysis. Data processing revealed an inherently dynamic NatA interaction with the ribosome, where the heterodimer adopts a multitude of conformations, which severely limits the local resolution **(Supplementary Figure 1).** Nonetheless, this preliminary structure revealed that, compared to yeast NatE^14^, human NatA accommodates a completely different non-intrusive off-center position near the PTE, in the following denoted as ‘distal’ site, independent of a nascent chain substrate or other RAFs.

Recently, we reported several structures of MAP2 on the ribosome^12^. The central placement of the enzyme on top of the PTE would not result in steric clash with NatA at its distal position, and might therefore allow a coincident occupation of the ribosome by these two sequentially active enzymes. As an attempt to stabilize NatA in the distal site, we added MAP2 to the complex and recorded a new cryo-EM dataset **(Supplementary Figure 2, Supplementary Table 1).**

Indeed, concurrent binding of human MAP2 constrained the conformational freedom of NatA and improved the local resolution to ∼3 Å, without interfering with the distal position of NatA **(Supplementary Figure 2)**. Therefore, this dataset was used for a detailed analysis of the NatA-80S interaction.

Consistent with the reported functions of Naa15, the two major contacts formed between NatA and the ribosomal surface are mediated by this adapter protein (interface of 1,100 Å^2^ calculated using PISA^35^). The first contact is formed by the longest α-helix of Naa15 (helix α34, residues 591-628) **(Supplemental Figure 3)**, which wedges in the widened major groove of the most exposed kink-loop of expansion segment ES7L(A) **(Figure 1a)** (proximal and extended: nucleotides 467-471 and 683-688) and bridges a large gap of over 55 Å to ES44L **(Figure 1b)**. The second contact to the PTE is formed by the N-terminal TPR motif (helices α1 and α2), which is positioned in between exposed ribosomal 28S rRNA helices (H19 and H24) and the peripheral lining of uL24 **(Figure 1c)**. 3D classification revealed that NatA can still assume a range of conformations despite the presence of MAP2 **(Supplementary Figure 4)**. The long flexible linkers of helix α34 (16 N-terminal residues: 573-588, and 24 C-terminal residues: 629-652) that connect the helix to the rest of Naa15 **(Figure 1)**, allow NatA to rappel further down towards the PTE while retaining all its ribosomal contacts.

**Figure 1:**
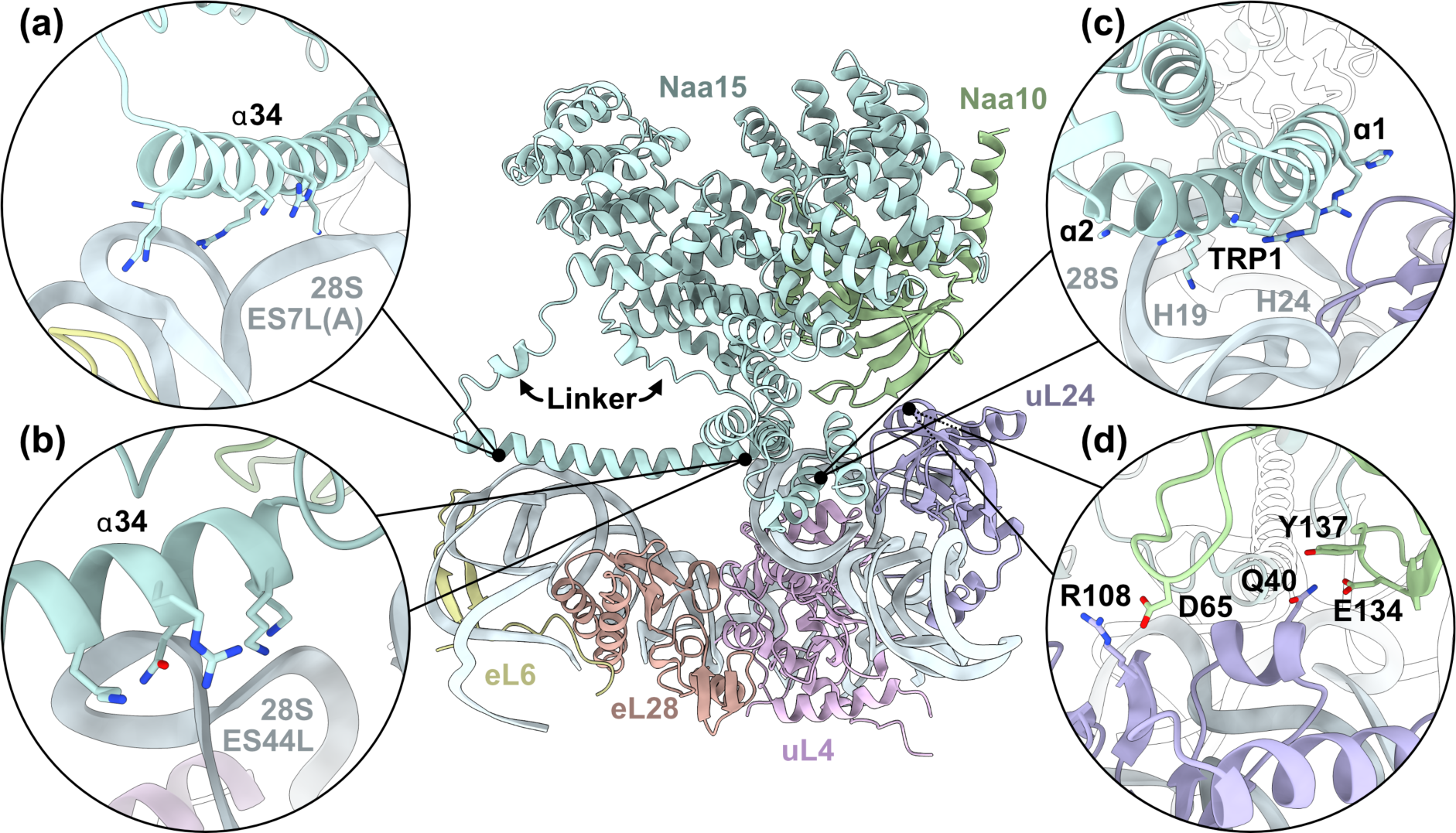
Structure of NatA in the distal site. Figure is made from the ternary NatA-MAP2-80S structure. Helix α34 anchors the NatA complex via electrostatic interactions with the phosphate backbone of **(a)** ES7L(A) and **(b)** ES44L. **(c)** The first two helices of Naa15 (α1 and α2) touch down on top of H19 and H24 in proximity to uL24. **(d)** In addition, Naa10 engages in multiple interactions with uL24.

While Naa15 contributes most prominently to the ribosome interactions, Naa10 also contacts uL24 in some of the NatA states that we captured **(Figure 1d)** (interface less then 100 Å^2^ calculated using PISA^35^). These interactions contain salt bridges (e.g. D65^Naa10^-R108^uL24^) and polar contacts (E134/Y137^Naa10^-Q40^uL24^). However, these contacts appear transient as NatA rotates in the distal site.

Compared to the crystal structures of NatA, the Naa15 solenoid undergoes rearrangements in the first two TPRs, when adapting to the distal site. Similar rearrangements have also been observed through the stabilizing interaction with HypK^17,18^. Of note, the bending of the first two Naa15 TPRs occurs in opposite directions upon HypK or ribosome binding **(Supplementary Figure 5)**.

While the same structural features of Naa15 are also employed in ribosome binding of *Sc*NatE^14^, *Hs*NatA uses the two contact sites to bind in a completely different position far off-center from the PTE, denoted here as the distal site **(Supplementary Figure 6)**. Without the steric constraint formed by Naa50 in the *Sc*NatE structure, *Hs*NatA binding becomes even more dynamic. When superimposing the structure of *Hs*NatE^18^ onto the distal position of *Hs*NatA on the ribosome, Naa50 is pointing far away from the tunnel exit. In this position it is completely solvent exposed and would not contact the ribosomal surface **(Supplemental Figure 7)**, indicating that human NatE might have a different binding site than NatA.

### MAP2 binding in presence of NatA

Up to 85% of all human proteins are N-acetylated and NatA carries out the majority of these modifications^3,36^. Unlike NatB or NatC NatA requires prior NME by methionine aminopeptidases^3,36^. Removal of the initiator methionine can hereby be carried out by MAP1 or MAP2 in eukaryotes.

Recently, we reported several structures of MAP2 on the ribosome^12^. The metalloprotease has a typical pita-bread fold with a MAP2-specific insertion domain that locates centrally on top of the PTE. In this position, MAP2 forms a vestibule-like structure to expose the active site towards potential emerging nascent chain substrates^12^.

In the NatA-MAP2-80S complex, MAP2 and NatA binding is not altered in the presence of the other factor and there is no direct interaction between them (at least their structured domains). **(Figure 2a)**. As described in detail before^12,37^, the strong binding of MAP2 on top of the PTE and the adaptable rRNA helix 59 is mainly due to the MAP2-specific insertion domain **(Figure 2a)** not present in the MAP1 family **(Supplementary Figure 3)**. In our previous cryo-EM structure of the human MAP2-80S complex, the long ES27L was recruited on top of the PTE by the N-terminus of MAP2 in a subset of particles (ES27L_out_ position). Now, the structure of the MAP2-NatA-80S complex shows ES27L locked away in the ES27L_in_ position at the 60S/40S interface (as also observed in our NatA-80S complex) **(Supplemental Figure 8)**. The absence of the MAP2-ES27L contact does not influence the MAP2 position on the PTE.

**Figure 2:**
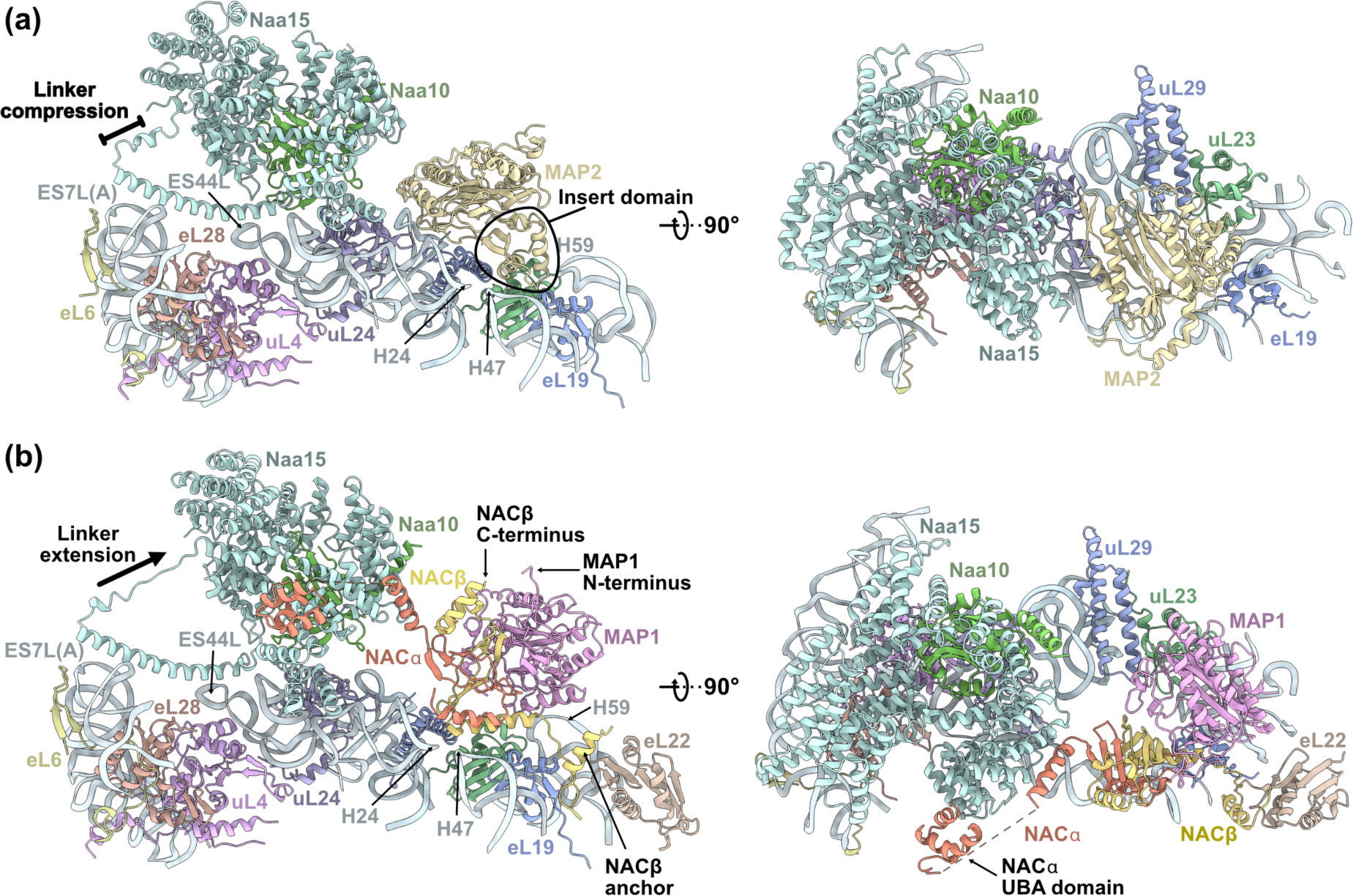
NatA forms coordinated assemblies with MAPs on the ribosome. **(a)** MAP2 is positioned centrally on the PTE. This interaction is mediated by the insert domain which binds on top of H59. The Naa15 linker is compressed into a helix and NatA is rotated further away from the PTE. **(b)** NAC mediates the interaction between NatA and MAP1. NAC uses its N-terminal NACβ anchor to contact eL19 and eL22. The C-terminus of NACβ points towards the N-terminus of MAP1. The UBA domain of NACα contacts the N-terminal Naa15 helices (α4-6). Overall, the NatA complex is rotated further down towards the PTE and the flexible linker is more extended.

### NAC mediates the interaction between NatA and Map1

The ternary NatA-MAP2-80S assembly reveals one possible way of coordinating NME with NTA on the ribosomal PTE. In eukaryotes, NatA substrates could alternatively be produced through the function of MAP1, instead of MAP2. Aside from the lacking insert domain in MAP1, the two enzymes mainly differ in the architecture of their N-terminal extension^10^. While the MAP2 N-terminus is unstructured and comprises segmented regions of alternatingly charged residues, the MAP1 extension harbours zinc-finger motifs and is implicated in NACβ binding^11^. The active center of both enzymes is highly conserved and their substrate specificity only shows small differences^38^.

To assess whether NatA and MAP1 can also bind together on the ribosome in preparation for successive NME and NTA, we mixed recombinant protein with purified human ribosomes for SPA cryo-EM analysis. After extensive local classification of a preliminary dataset in cryoSPARC, we identified MAP1 and NatA on the ribosome in complex with endogenous NAC, which had co-purified with the 80S monosomes despite the high-salt condition (500 mM KOAc) during gradient centrifugation. To enrich NAC on the ribosome, we cloned, expressed and purified NAC from insect cells and reconstituted the quaternary NatA-NAC-MAP1-80S assembly for cryo-EM data collection **(Supplementary Figure 9, Supplementary Table 1)**.

Several rounds of focused 3D classification were required to deal with the extensive heterogeneity that result from this mix of factors on the ribosome **(Supplementary Figure 9)**. NatA, NAC and MAP1 all exhibit continuous flexibility and explore a range of different conformations. Together, these factors assemble into a dynamic ring-like arrangement that engulfs the tunnel exit, leaving a large lateral gap towards uL29 **(Figure 2b)**. Classification down to a small subset of particles (∼4 %) revealed the quaternary assembly in a more stabilized configuration, allowing a detailed analysis of the molecular interactions. Sandwiched in between NatA and MAP1, NAC is positioned on top of 28S rRNA helices H24 and H47, in the same position where has been found in the absence of other factors **(Figure 2b)**^22^. Alike NatA, NAC does not require a nascent chain to bind the ribosome and to position itself at the PTE. NAC forms a peculiar heterodimer (α and β subunit of 215 and 206 residues, respectively) that assembles on a β-barrel-like core, flanked by two α-helical pairs and long flexible hydra-like termini. Situated at the PTE, the NAC dimerization domain packs tightly against the lateral side of MAP1 with the β-barrel-core placed almost perpendicular to the inner MAP1 cavity, without blocking substrate entry to the active site **(Figure 2b)**.

The long terminal extensions of NAC can act over longer distances to facilitate various functions^11,22,39–41^. When NAC is positioned upright at the PTE, the N-termini of NACα and NACβ point towards the ribosome. The N-terminus of NACβ is resolved up to the short anchoring helix that facilitates NAC recruitment to the ribosome by binding next to eL22 and eL19^40,41^ **(Figure 2b)**. The NACα N-terminus is not visible in our reconstruction, but points towards the Naa15 interface. A potential role of this extension in the regulation of ribosome binding has previously been reported^27,39^. The unstructured C-terminal extension of NACβ has been shown to mediate MAP1 recruitment via a conserved hydrophobic patch that binds the N-terminal extension of the aminopeptidase^11^. While this interaction is not visible in our reconstruction, the trajectory of the NACβ C-terminus clearly points towards the N-terminus of MAP1 **(Figure 2b)**. Despite the presence of NatA and the absence of nascent chains, NAC and MAP1 can bind to the PTE in the same way as previously described in the presence of a nascent chain substrate^11^.

In contrast to the unstructured NACβ extension, NACα includes an additional UBA-like domain at the very C-terminus **(Supplementary Figure 3)**. UBA-like domains are protein-protein interaction modules folded in a three-helix bundle and are found in e.g. the ubiquitin/proteasome pathway^42^. In the presence of NatA, NACα-UBA touches down on the Naa15 N-terminal helices (helices α4, α5 and α6) **(Figure 2b)**. The NACα-UBA/Naa15 interface involves a hydrophobic core surrounded by polar and ionic interactions **(Supplemental Figure 10)**. In context of a nascent chain ‘handover’ from NAC to the signal recognition particle (SRP), responsible for co-translational protein targeting to the ER membrane, the NAC UBA domain was found laterally attached to the SRP key-player SRP54^22^ **(Supplementary Figure 10)**.

In complex with NatA, the flanking NACα helix that connects the β-barrel core to the UBA domain (denoted here as NACα contact helix (NACα−cH)) **(Supplemental Figure 3)** aligns outside Naa15 TPR4-TPR5 (helices α8, α9 and α10). This interaction includes a central salt bridge (K142^NACα^-E173^Naa15^) and an aromatic cluster around F143^NACα^, and is complemented by various peripheral polar contacts **(Supplementary Figure 11).** Together, the NACα-cH and NACα-UBA share an interface of almost 1000 Å^2^ with Naa15 (calculated using PISA^35^)). Of note, the NatA modulator HypK displays homology to NACα and also contains a UBA domain that is preceded by an α-helix which employs this exact interface with Naa15 TPR4-TPR6^16^ **(Supplementary Figures 3 and 11)**. For NACα however, the connecting cH helix is much shorter than the corresponding HypK helix. This structural difference apparently contributes to a change in the UBA placement at the Naa15 solenoid. While the NACα-UBA contacts the N-terminal TPR2 and TPR3, HypK-UBA is positioned diametrically opposed and in contact with the Naa15 C-terminal TPRs **(Supplementary Figure 11)**. The function of the Naa15 solenoid as a binding hub for external helical elements is even further corroborated by our finding that the Naa10 C-terminal helix also crawls along Naa15 TPR6-TPR7, in a very similar fashion as the NACα-cH **(Supplementary Figures 3 and 12)**.

Overall, the quaternary assembly of NatA, NAC, MAP1 with the ribosome is highly dynamic. Despite the numerous contacts that NatA forms to the ribosome and to NACα, it is still free to rotate in the distal site while retaining all of its contacts **(Supplementary Figure 13)**. Compared to our ternary NatA-MAP2-80S structure, the NatA complex is rotated 25° down towards the PTE and the first TPRs are shifted further away from uL24 **(Supplementary Figure 14)**. Throughout this motion, the N-terminal liker (residue 571-588) can extend and compress between 38 Å and 50 Å to allow the dynamic rotation of Naa15 **(Supplementary Figure 14)**.

## Discussion

Two MAPs, two NMTs and five NATs need to be coordinated at the crowded PTE in the context of a rapidly translating ribosome^2^. Once the nascent chain starts to emerge from the ribosome, the first two residues can become targets for modification by this elaborate pool of factors. While the nascent chain is still short and barely protruding from the PTE, these target residues are still in a predictable and accessible position. To exploit this brief window of opportunity, the enzymatic processes that mature a growing nascent chain need to be well coordinated. NME by MAP1 or MAP2 and subsequent NTA by NatA are the most widely executed co-translational modifications in human cells^3^. Given the relative underrepresentation of these enzymes compared to the ribosome, their binding, activity and eventual release must be tightly controlled.

Here, we identified that NatA can bind the ribosome in a non-intrusive off-center distal site. While ribosome binding of many RAFs is mutually exclusive, the NatA distal site would not interfere with binding of most other known RAFs at the PTE, namely MAP1, MAP2, Ebp1, NatB, NAC, RAC, SRP and SEC61 **(Figure 3a)**. A coincident binding of NatA with factors that function up- or downstream of acetylation, might allow for a more coordinated handover of the nascent chain substrate in between the subsequent maturation steps. To explore two different routes that might enable the coordinated handover between NME and NTA, we determined two cryo-EM structures of NatA in complex with either MAP2 or MAP1.

**Figure 3:**
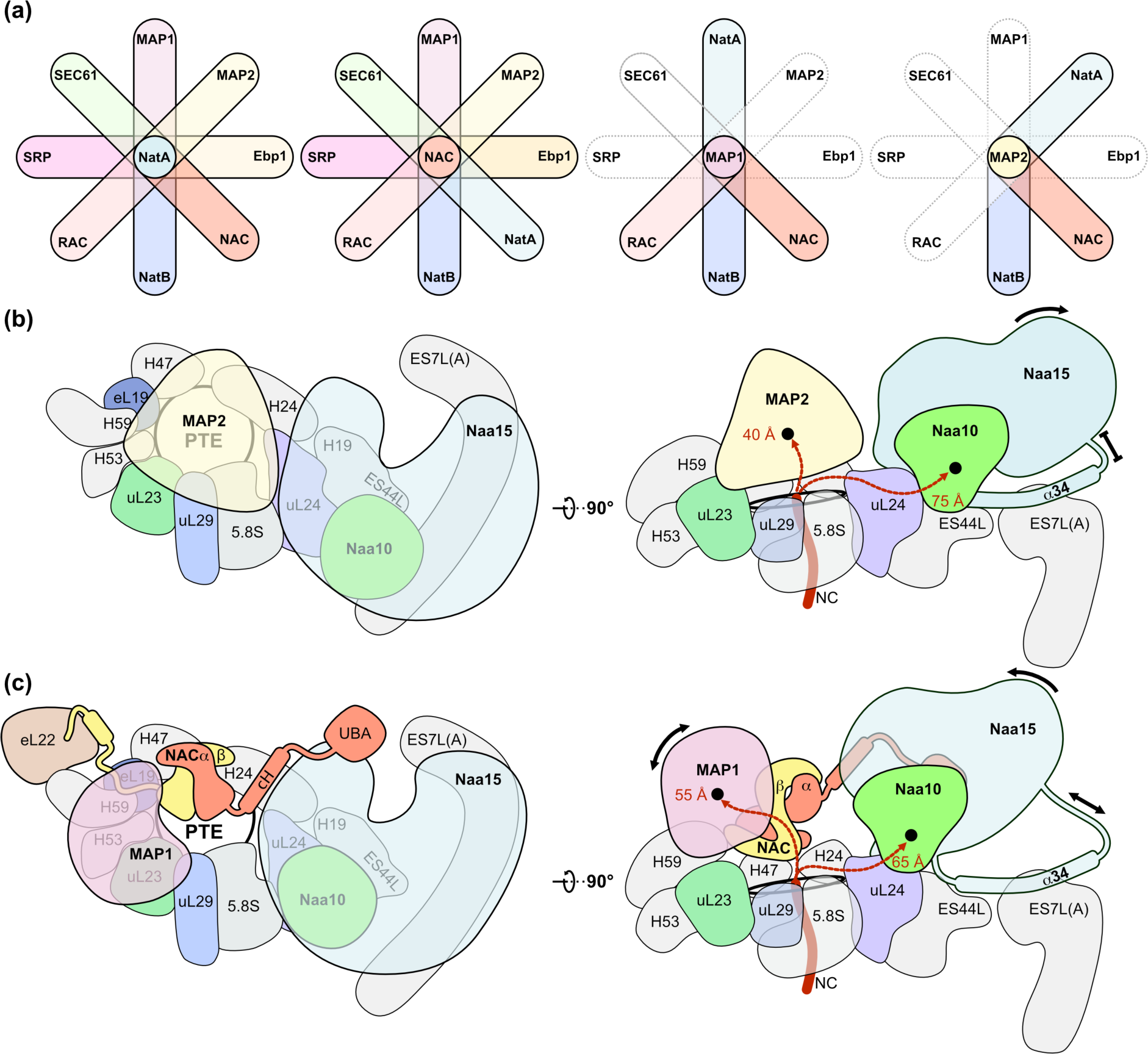
Coordinated multi-factor assemblies compile on the PTE. **(a)** Diagrams show the different RAF combinations that could form on the ribosome without clashing, including MAP1 (this study), MAP2^12^, NatA (this study), Ebp1^37^, NAC (this study), NatB^43^, RAC^44^, SRP^22^ and SEC61^45^. The off-center distal position of NatA and the non-intrusive NACβ anchoring position does not interfere with binding of any of these factors. In contrast, MAP1 would clash with MAP2, Ebp1, SRP and SEC61, while MAP2 would clash with MAP1, Ebp1, RAC, SRP and SEC61. **(b)** Schematic top and side view of the ternary NatA-MAP2-80S assembly. MAP2 is positioned centrally on the PTE and does not interact directly with NatA. NatA is anchored to ES7L(A) and ES44L with its α34 helix and the connecting linker is compressed. In this assembly, the distance to the active sites of MAP2 and Naa10 are 40 Å and 75 Å (measured from 28S rRNA G2416^46^). **(c)** Schematic top and side view of the quaternary NatA-NAC-MAP1-80S assembly. NAC is anchored with its NACβ helix adjacent to eL22 while the dimerization domain is placed on top of H47 and H24. The NACα-cH and UBA domain contact Naa15 and the dimerization domain is in close contact to MAP1. The NatA complex is still anchored on ES7L(A) and ES44L with its α34 helix, but the connecting linkers are almost completely extended. The distance to the active site of MAP1 and Naa10 would be 55 Å and 65 Å for the two enzymes, respectably (measured from 28S rRNA G2416^46^). Both NatA and MAP1 remain dynamic within the assembly and can rotate towards and away from the PTE.

In the NatA-MAP2-80S structure, the individual binding sites of MAP2 and NatA were not affected by the presence of the other factor **(Figure 3b)**. Only the rotational freedom of NatA was limited, resulting in a smaller angle of rotation due to the PTE occupation of MAP2, which allowed for a higher-resolution reconstruction of NatA in the distal site. In our previous cryo-EM structures of the MAP2-80S complex, we reported that the enzyme can undergo a dynamic rotation around the insert domain in the presence of nascent chains^12^. Such a rotation of MAP2 might allow NatA to rappel further down towards the PTE and position the Naa10 subunit closer to the emerging nascent chain. The motion of NatA towards MAP2 might also cause the displacement of the latter, to make room for binding of the subsequent processing factor. Within the ternary NatA-MAP2-80S assembly, an emerging nascent chain would need to bridge a gap of ∼40 Å and ∼75 Å to reach the active sites of MAP2 and Naa10, respectively (measured from 28S rRNA G2416^46^) **(Figure 3b)**.

While human MAP2 binds rigidly at the PTE in the absence of nascent chains and other factors^12^, MAP1 binding to the ribosome in complex with NAC is inherently dynamic. In the quaternary NatA-NAC-MAP1-80S assembly, NAC employs its hydra-like structure to coordinate the two successively active enzymes at the PTE. Despite the multitude of interactions between each factor and the ribosome, as well as between the factors themselves, the entire ring-like assembly remains highly flexible. Especially NatA and MAP1 can rotate to and away from the PTE without losing contact to NAC and the ribosome **(Figure 3c)**. This structural adaptability might be especially important to accommodate different nascent chains with varying sequence and properties that emerge from the PTE during translation. The ability to adapt but remain PTE associated when the nascent chain emerges could give these factors more time to carry out NME and NTA before making room for downstream processing factors or chaperones. Compared to the NatA-MAP2-80S complex, the indirect interaction between MAP1 and NatA, mediated by NAC, results in a shorter distance of the Naa10 active site to emerging substrates. Measured from G2416 of the 28S rRNA^46^, nascent chains would need to extend for 55 Å and 65 Å to reach the active sites of MAP1 and Naa10, respectively **(Figure 3c)**.

Via the central placement on top of H24 and H47 in between NatA and Map1, NAC shields off one side of the PTE. In this configuration, the lateral wall shaped by the NAC dimerization domain might aid in guiding emerging nascent chain substrates more efficiently into the active site of MAP1 **(Figure 3c)**. In contrast, the vestibule-like structure of MAP2 leaves an opening only towards H24 and H47, and completely shields off the opposing direction towards uL29.

While the interaction between the NACβ C-terminus and MAP1 is not visible in our reconstruction, the interaction between NACα and NatA is well defined. The parallel placement of the NACα-cH helix alongside helices α8-10 could also be observed in the NatA-HypK complex. To date, the precise function of HypK is not understood. In our structures, we observe that structural adaptations in the first two TPR repeats of Naa15 (α1-4) occur when NatA adopts the distal position on the ribosome **(Supplementary Figure 5)**. Interestingly, HypK binding also induces a similar bending of these helices, however in opposing direction^16,17^. The stabilizing effect of HypK binding on NatA has been well described^16,17^. Perhaps the rigidification of the Naa15 scaffold by HypK binding would prevent the distal site interaction and restrict its structural plasticity, unless HypK rearranges in some way. These observations would corroborate the proposed function of HypK as a NatA inhibitor. Reports that HypK contacts nascent chains and co-localizes with polysomes contradicts the inhibitory function on ribosome binding^15,19^. NatA has the broadest substrate specificity of any N-Acetyl Transferase^3^ and could potentially co-occupy the PTE with most other RAFs known to date. Furthermore, the Naa15 scaffold constitutes a central binding hub for a multitude of factors (e.g. Naa10, Naa50, HypK, NACα). This plethora of possible Naa15 interactions needs to be regulated in some way. Perhaps HypK acts as a ‘specificity filter’ that ensures that NatA is only employed on the ribosome in the right condition, possibly in the presence of other factors that can induce rearrangements in HypK. Herein, NAC would be a prime candidate, as its NACα−cH utilizes the same interaction surface on Naa15 as the HypK-cH. Given the low nanomolar affinity of human HypK to the NatA complex^16,17^ it has been suggested that in human cells, these proteins form an obligate ternary complex *in vivo*^16^. In our NatA-MAP2-80S structure, we observed no direct interaction between NatA and MAP2. If and how HypK might be displaced in the presence of MAP2 remains to be explored.

The diverse functions inherent to the terminal extensions of NAC are only partially understood. The precise function of the ∼70 residue long N-terminal NACα extension remains to be described. Interestingly, the ‘QQAQLAAAE’ motif in the NACα-cH which mediates the Naa15 interaction described here is duplicated in the N-terminal extension of NACα (‘QQAQLAAAAE’). Whether the N-terminus of NACα can also contact Naa15 remains to be explored.

Connected to the NACα-cH via a short unstructured linker, NAC possesses a UBA-like domain that touches down on the N-terminal helices of Naa15. In complex with SRP, the NAC UBA domain was found to also contact SRP54^22^. This interaction is highly similar to the NACα-Naa15 interaction, as the UBA domain is also positioned on two parallel helices with its exposed hydrophobic patch **(Supplementary Figure 10)**. Most SRP targets are not subject to NME and NTA by MAP1 and NatA while aberrant acetylation of such nascent chains has been shown to inhibit translocation^47^.

The coordination of NME and NTA by NAC or the coordination of SRP recruitment by NAC are therefore likely two mutually exclusive processes.

Most RAF interactions with the ribosome can occur independent of a nascent chain substrate (e.g. MAP2^12^, Ebp1^37^, NAC, NatA, NatB^43^). Furthermore, all RAF enzymes only target the first two residues of the nascent chain. These residues will be most accessible while nascent chains are still short (e.g. <50-60 residues). Given the high speed of translation, the flexibility and variability of the nascent chain, there would likely not be enough time to wait for the N-terminal residues to emerge from the PTE before the enzymes associate to the ribosome. Instead, we postulate that muti-protein ‘starter-kits’ form on the PTE to reside in a reactive state, and await the emerging substrate. Once the nascent chain starts to grow beyond the PTE, it might induce conformational changes in these RAFs (e.g. a rotation of MAP2^12^) that allow the recruitment of downstream factors. A ‘starter-kit’ composed of MAP1, NatA and NAC would combine the functionalities of NME, NTA and molecular chaperoning. Perhaps, one alternative ‘starter-kit’ might compose of RAC, NatA and MAP1. A major question that remains is how the composition of these start assemblies is controlled.

## Methods

### Sample preparation

Genes encoding NACα, NACβ and Naa10 were purchased from IDT peptide. Genes encoding Naa15 and MAP1 were subcloned from vectors pFastBac-HIS-HsNaa15 and pET24d-HIS-TEV-HsMAP1, respectively.

NatA was cloned into pFastBacDuet plasmid with Naa10 expressed under the polyhendrin promotor with a 3C-protease cleavable C-terminal HIS-tag (Naa10-GSGS-3C-GSGS-10HIS). Untagged Naa15 was co-expressed under the p10 promotor. N-terminally HIS-tagged MAP1 was expressed under the polyhendrin promotor from pFastBacDuet plasmid with a 3C-protease recognition site between the HIS-tag and MAP1 (10HIS-GSGS-3C-GSGS-MAP1). NAC was also cloned into pFastBacDuet plasmid with NACα expressed under the polyhendrin promotor with an N-terminal 3C-cleavable Step-Tag II (Step-Tag II-GSGS-3C-GSGS-NACα). NACβ was co-expressed from the p10 promotor without an affinity tag.

All proteins were expressed in Sf9 cells. Briefly, insect cells were grown at 27°C and 80 rpm to be infected at a density of 2 × 10^6^ cells/ml. Expression was continued for 72h and cells were harvested at by centrifugation at 1500 × g. Pellet was washed once with PBS, flash frozen in liquid nitrogen and stored at -80°C until further use. For purification, pellets were lyzed in microfluidizer in the presence of 1 × protease inhibitor (Roche) and cleared by ultracentrifugation at 50,000 × g for 20 min. Cleared lysate was passed through 0.2 µm syringe filter and proteins were purified at 4°C. The purification of MAP1, NAC and NatA were done by a two-step purification scheme. To produce pure NAC, *StrepII-GSGS-3C-GSGS-NACα* and *NACβ* were co-expressed in Sf9 cells. Recombinant protein was captured on StepTactin™ Agarose (Purecube). To obtain pure NatA, *Naa10-GSGS-3C-GSGS-10H* and *Naa15* were co-expressed in insect cells. Naa10 was captured on Ni^2+^-Agarose beads (Qiagen). *10H-GSGS-3C-GSGS-MAP1* was also purified on Ni^2+^-Agarose beads. After the capture step, NAC, NatA and MAP1 purification were done by the same protocol. Affinity tags were cleaved-off with 3C protease and eluted protein was concentrated to 10 mg/ml. To obtain stoichiometric complexes and pure MAP1, samples were subjected to SEC on a S200 16/600 (Cytiva) column in SEC buffer (20 mM HEPES KOH, 150 mM KOAc, 5 mM MgOAc_2_, 1 mM TCEP, pH 7.4). Purity of proteins was evaluated by SDS-PAGE and analytical SEC on S200 increase 3.2/300 GL (Cytiva) **(Supplementary Figure 15)**. *Hs*MAP2 was expressed in Sf9 cells, as previously described^12^.

Human 80S ribosomes were isolated from HeLa S3 cells as described^12,37^, with the sucrose cushion adjusted to a KOAc concentration of 500 mM. For cryo-EM analysis of the NatA-80S complex, ribosomes were subjected to SEC (20 mM HEPES KOH, 600 mM KOAc, 5 mM MgOAc_2_, 1 mM TCEP, pH 7.4) on a Superose 6 10/300 column (Cytiva) to remove any co-purified RAFs from the PTE. Afterwards, the buffer was adjusted to 20 mM HEPES KOH, 150 mM KOAc, 5 mM MgOAc_2_, 1 mM TCEP, pH 7.4. For cryo-EM datasets of NatA-NAC-MAP1-80S, ribosomes were not subjected to high-salt SEC to additionally retain endogenous co-purified NAC. After purification, ribosomes were flash frozen in liquid nitrogen and stored at -80°C until further use.

### Cryo-EM grid preparation

Before sample vitrification, copper grids were glow-discharged in the Solarus plasma cleaner (Gatan, Inc.) for 60 sec in oxygen atmosphere. For the data acquisition of NatA, 500 nM of high-salt washed 80S ribosomes were mixed with 12.67 µM of purified NatA. For the complex of MAP2 and NatA, 500 nM of high-salt washed 80S ribosomes were incubated with 30 µM of MAP2 and 30 µM of NatA for 30 min at room temperature. The sample preparation for the NatA-NAC-MAP1-80S complex was performed at 37°C. Here, 500 nM of 80S ribosomes (not high-salt cleaned) were incubated with 5 µM of NAC for 10 min at 37°C. NatA and MAP1 were added one after the other, with 10 min incubations in between, each to a final concentration of 15 µM. From each sample, 3 µl were frozen in liquid ethane with the Vitrobot Mark IV (FEI company) on 2/1 Quantifoil Multi A holey carbon supported grids (Quantifoil, Multi A, 400 mesh). Freezing was carried out with Whatman #1 filter papers at 4°C with a blot force of 0, 10 sec wait time and 100% humidity. Grids were stored in liquid nitrogen until data acquisition.

### Data collection

Two datasets of the NatA-80S sample were collected on a Glacios transmission electron microscope (Thermo Fisher Scientific) operated at 200 keV with the Falcon 3 direct electron detector (Thermo Fisher Scientific) that collected at a pixel size of 1.223 Å/pixel and a magnification of 120,000. The total dose per was 51.65 and 55.07 e^-^/Å^2^ for the two datasets, respectively. Three datasets of the NatA-NAC-MAP1-80S assembly were also collected on the Glacios transmission electron microscope at a magnification of 120,000 and a pixel size of 1.223 Å/pixel. The total dose for the three datasets was 53.50, 53.97 and 53.96 e^-^/Å^2^ respectively. The dataset of NatA-MAP2-80S was collected at ESRF on a Titan Krios electron microscope (Thermo Fisher Scientific, FEI Company) on a K3 Summit direct electron detector (Gatan, Inc.) that collected movies at a pixel size of 0.84 Å/pixel at a magnification of 105,000. The total dose of the NatA-MAP2-80S dataset was 41.28 e^-^/Å^2^. The data acquisitions were set up and monitored with EPU (Thermo Fisher Scientific).

### Data processing

Detailed descriptions of the processing schemes for all datasets can be found in the supplementary information. Briefly, movies were imported and pre-processed in CryoSPARC^48^, including Patch Motion Correction and Patch CTF Estimation. Ribosomes were picked with the Blob Picker and extracted for 2-3 subsequent rounds of 2D classification. Selected particles were subjected to ab-initio reconstruction into 3 classes, followed by Heterogeneous refinement. Classes containing 80S ribosomes were pooled and refined using the Homogenous Refinement job in CryoSPARC. Remaining particles were subjected to several rounds of focused 3D Classification with different masks to subsequently deal with the compositional and continuous heterogeneity of the factors at the PTE. After classification, masks were generated to subtract the 80S signal, with the Particle Subtraction job and local masks around the tunnel exit were used for final Local Refinements. Local Resolution Estimation jobs were run for global and local refinements.

### Cryo-EM model building, refinement, and analysis

The high-resolution cryo-EM structure of the human ribosome (PDB ID: 6ek0)^49^ was used as a starting point for model building. For building of NAC and NatA, Alphafold models were generated to also predict the flexible insertions and extensions^50^. For building of the ternary NatA-MAP2-80S complex, our MAP2-80S structure was used as a starting model (PDB ID: 8ONY)^12^. For building of the NatA-NAC-MAP1 interaction, the cryo-EM structure of NAC-MAP1-80S was used (PDB ID: 8P2K)^11^. For model building, the component proteins were first rigid body fitted into the cryo-EM map using UCSF Chimera-X^51^. Atomic model building was performed in Coot^52^ and the preliminary model was refined and validated in PHENIX suite^53,54^. The refinement statistics for all cryo-EM datasets are shown in Supplementary Table 1.

## Acknowledgements

We acknowledge excellent technical support by Marina Pelzl, Britta Klem and Astrid Hendricks. We acknowledge access to the infrastructure of the cryo-EM Network (HDCryoNET) at Heidelberg University, and support by Dirk Flemming (BZH), Jan Rheinberger (BZH), Lutz Nücker (BZH) and Götz Hofhaus (Bioquant). All cryo-EM grids and preliminary datasets, leading up to the final high-resolution datasets, were screened and acquired in our in-house facilities. We acknowledge the European Synchrotron Radiation Facility for provision of beam time on CM01 and we would like to thank Michael Hons for assistance. Further, we acknowledge the services SDS@hd and bwHPC supported by the Ministry of Science, Research and Arts Baden-Württemberg and the German Research Foundation through grants INST 35/1314-1 FUGG and INST 35/1134-1 FUGG. This work was supported by the Leibniz Programme of the Deutsche Forschungsgemeinschaft to I.S. (SI 586-6).

## Author contributions

M.A.K., K.W. and I.S. designed the study and wrote the paper. M.A.K. generated all DNA constructs, purified proteins and ribosomes. Cryo-EM grids were prepared by M.A.K. and data were acquired and processed by M.A.K.. K.W and M.A.K. build the atomic models and M.A.K., K.W. and I.S. analyzed the data.

