## Supplementary Information for "Multi-protein assemblies orchestrate co-translational enzymatic processing on the human ribosome"

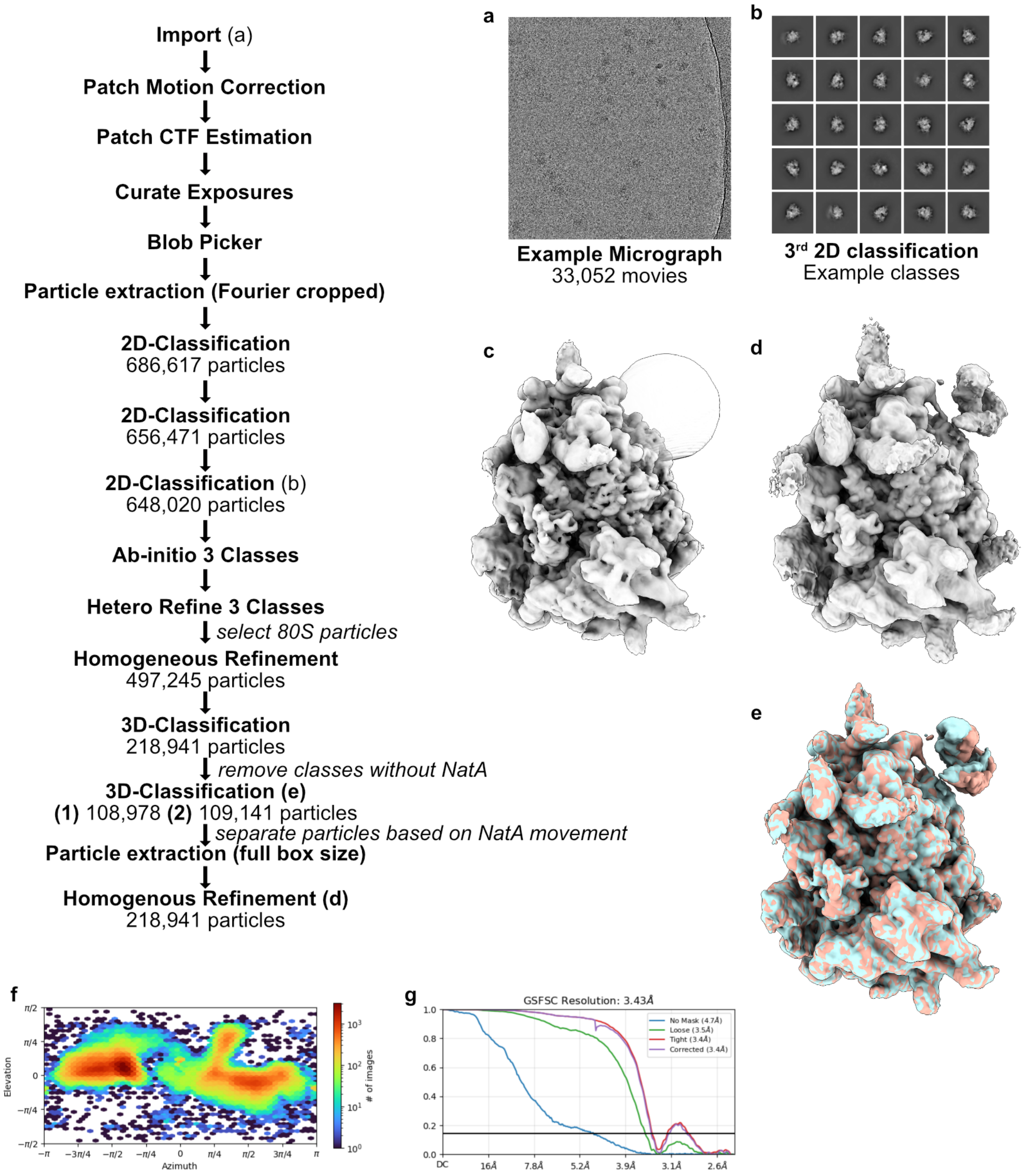


**Supplementary Figure 1: Cryo-EM data processing for the *Hs*NatA-80S sample. (a)** Two datasets were acquired to obtain 33,052 movies. **(b)** After pre-processing in CryoSPARC, extracted particles were subjected to three rounds of 2D-classification. Three ab-initio classes were generated from the remaining particles and used to seed a Heterogeneous refinement. 80S particles were selected and subjected to Homogenous Refinement. **(c)** A spherical mask was generated to encompass the distal site of NatA and used to initialize a 3D classification to remove particles without NatA. **(d)** Particles with NatA were subjected to Homogenous refinement. **(e)** To further subclassify particles based on the motion of NatA, a second 3D classification into two classes was performed. Both particle subsets were finally subjected to Homogenous refinement and superimposed to visualize the differences in NatA binding. **(f)** Angular distribution plot of particles used in the Homogenous refinement shown in (d). **(g)** FSC curves from the Homogenous refinement shown in (d).

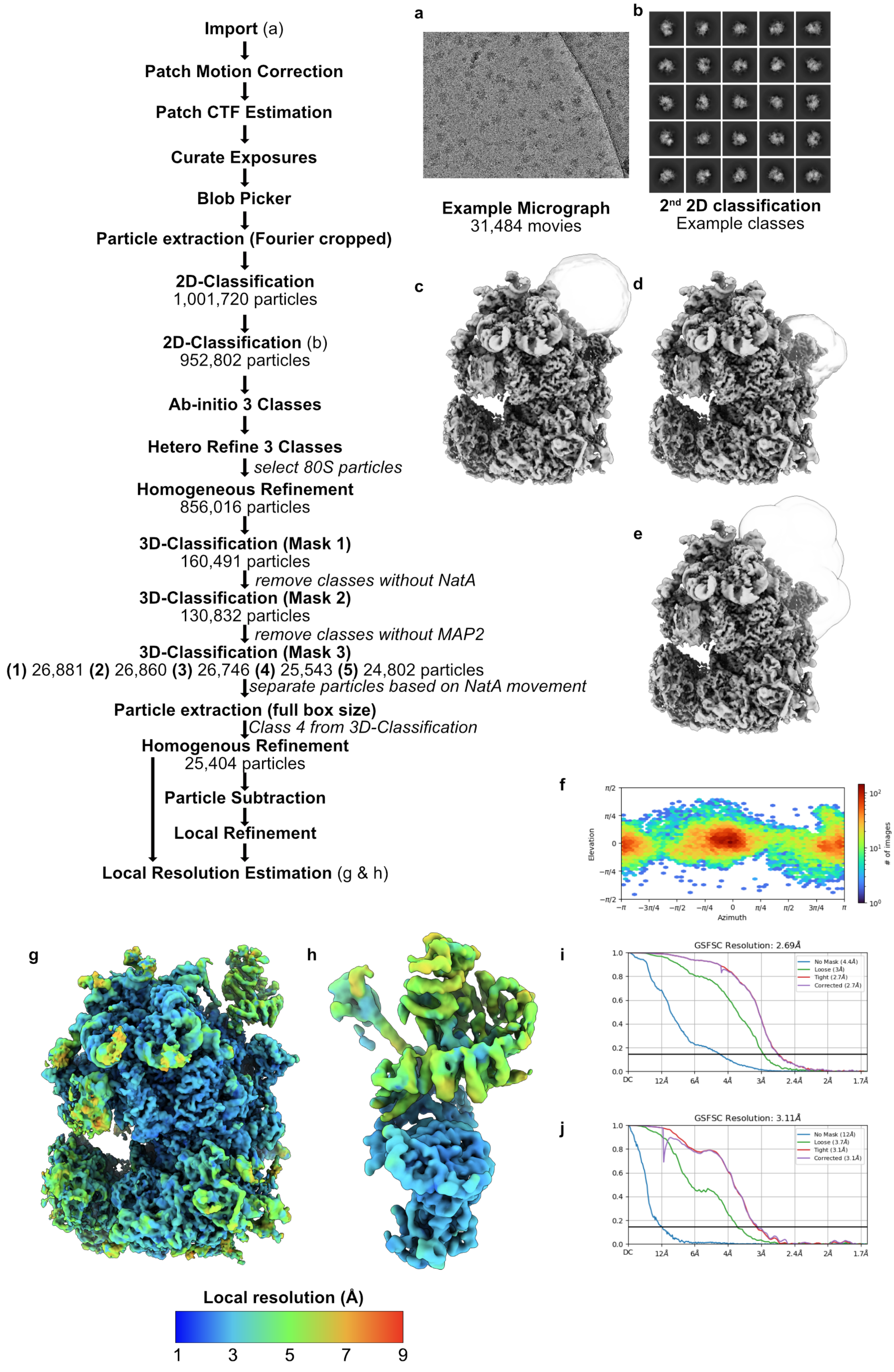


**Figure description is located on the next page**

**Supplementary Figure 2: Cryo-EM data processing for the *Hs*NatA-*Hs*MAP2-80S sample. (a)** A dataset was acquired to obtain 31,484 movies. **(b)** After pre-processing in CryoSPARC, extracted particles were subjected to two rounds of 2D-classification. Three ab-initio classes were generated from the remaining particles and used to seed a Heterogeneous refinement. 80S particles were selected and subjected to Homogenous Refinement. **(c)** A spherical mask was generated to encompass the distal site of NatA and used to initialize a 3D classification to remove particles without NatA. **(d)** A second mask was made to encompass the binding site of MAP2 and used to initialize a second 3D classification to remove particles without MAP2. **(e)** A larger mask was generated to encompass the binding site of NatA and MAP2 and used for a third 3D classification to sort particles based on the motion of NatA. One class yielded a higher local resolution around NatA. Corresponding particles were re-extracted without Fourier cropping and subjected to Homogenous refinement. **(f)** Angular distribution of particles used for the final Homogenous refinement. **(g)** Local resolution estimation of the final Homogenous refinement. **(h)** After Homogenous refinement, a mask was generated to subtract the 80S signal. The mask shown in (e) was subsequently used in a local refinement. Local resolution estimation was performed on the resulting map. **(i)** FSC curves for the Homogenous Refinement. **(j)** FSC curves for the Local Refinement.

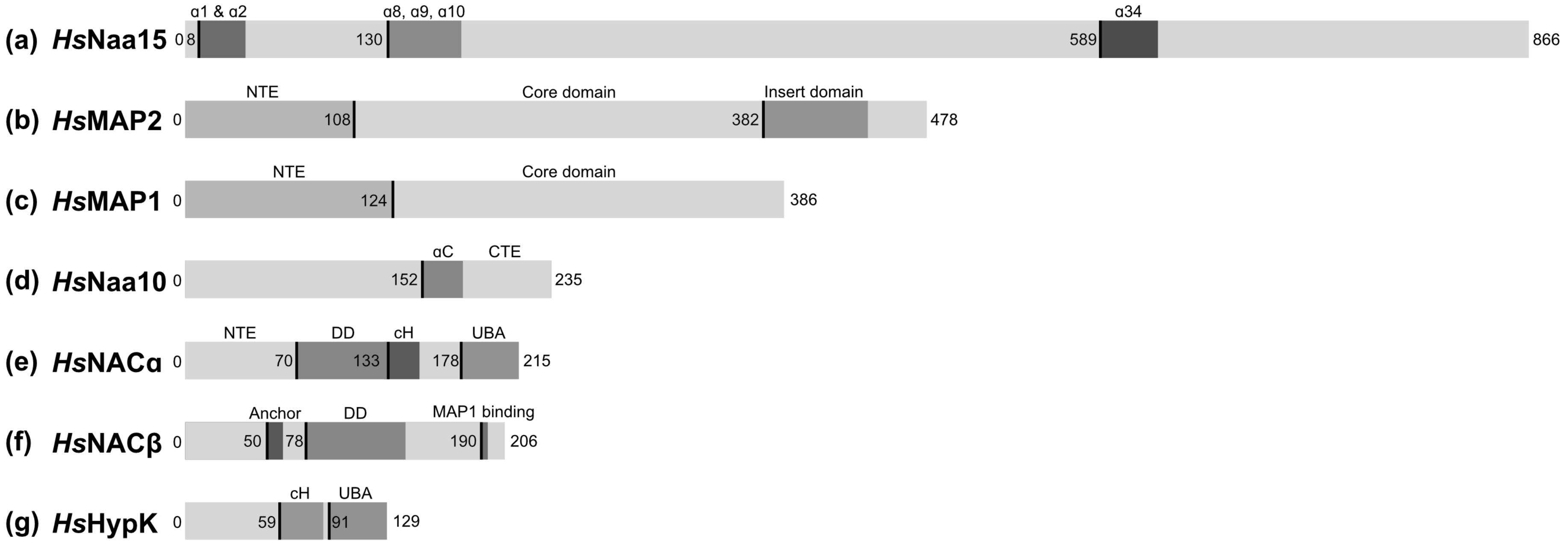


**Figure description is located on the next page**

**Supplementary Figure 3: Domain organization of ribosome associated factors. (a)** Domain organization of Naa15. The ribosomal contacts (helices $\alpha$1, $\alpha$2 and $\alpha$35) are highlighted, as well as the helices $\alpha$8, $\alpha$9 and $\alpha$10 which contact the NAC$\alpha$-cH and HypK-cH. **(b)** Domain organization of MAP2. The length of the unstructured N-terminal extension, as well as the position of the insert domain are indicated. **(c)** Domain organization of MAP1. The N-terminal extension (NTE) and core domain are highlighted. **(d)** Domain organization of Naa10. The C-terminal helix ($\alpha$C) and unstructured C-terminal extension are labelled. **(e)** Domain organization of NAC$\alpha$. The dimerization domain (DD) is located centrally within the protein sequence. Adjacent to the dimerization domain, the NAC$\alpha$-cH that contacts Naa15 is highlighted, as well as the three-helix bundle UBA domain. **(f)** Domain organization of NAC$\beta$. The dimerization domain (DD) is located centrally within the protein sequence. N-terminally, the helical anchor which mediates ribosome binding is highlighted. C-terminally, the conserved hydrophobic stretch that mediates MAP1 recruitment is highlighted[1]. **(g)** Domain organization of the NAC$\alpha$ homologue HypK. The HypK-cH that contacts Naa15 is highlighted, as well as the UBA domain.

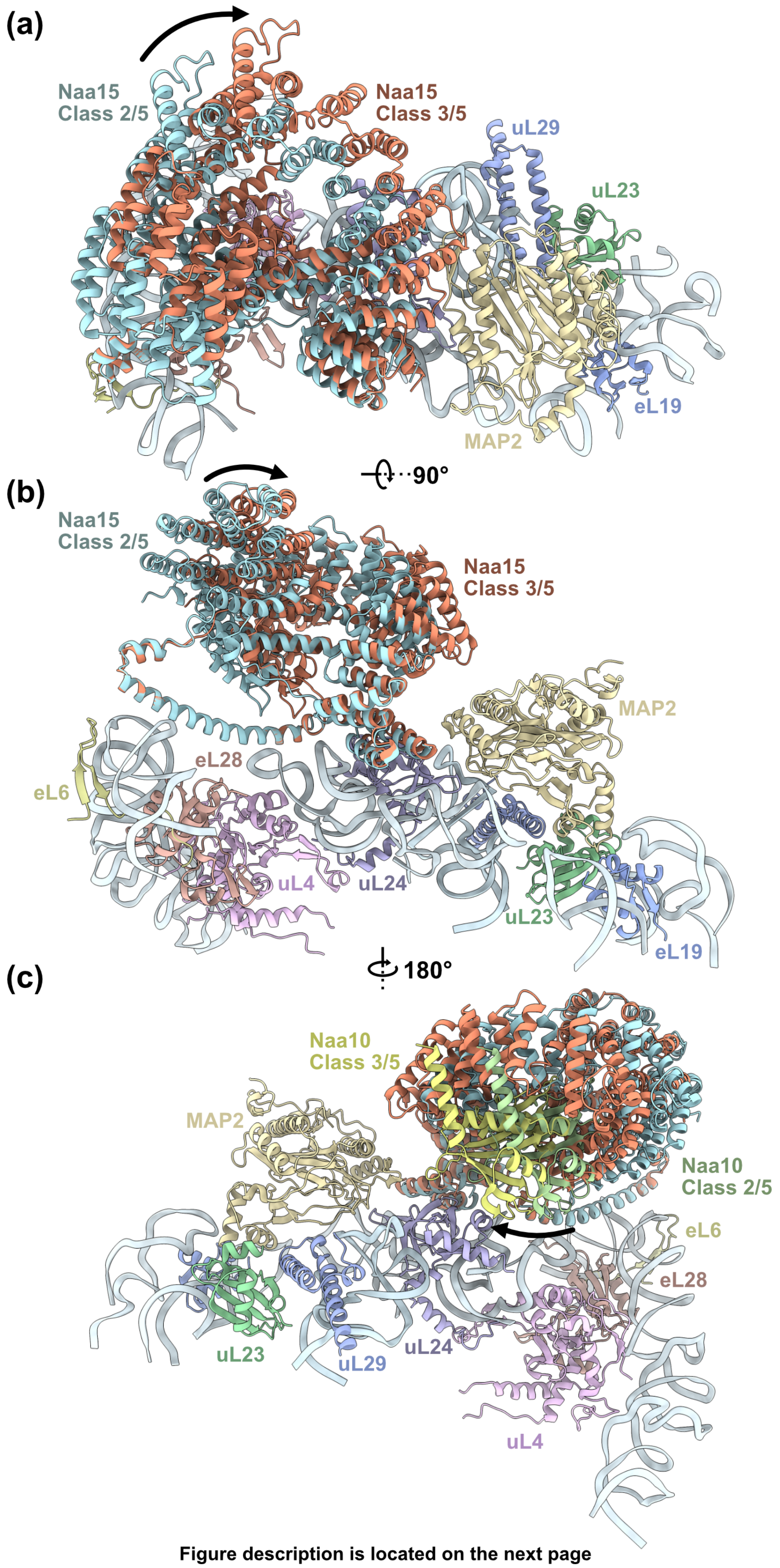


**Figure description is located on the next page**

**Supplementary Figure 4: Dynamics of NatA in the ternary NatA-MAP2-80S complex.** 3D classification revealed that NatA still moves dynamically in the distal site, despite the presence of MAP2. In total, five classes were obtained in the final 3D classification with slightly different modes of NatA binding **(Supplementary Figure 2)**. Classes 2 and 3 showed the strongest differences and are compared in this figure. **(a)** NatA twists and **(b)** rotates in the distal site but retains all contacts to the ribosome. **(c)** When NatA rotates towards the PTE, Naa10 is shifted along the surface of uL24.

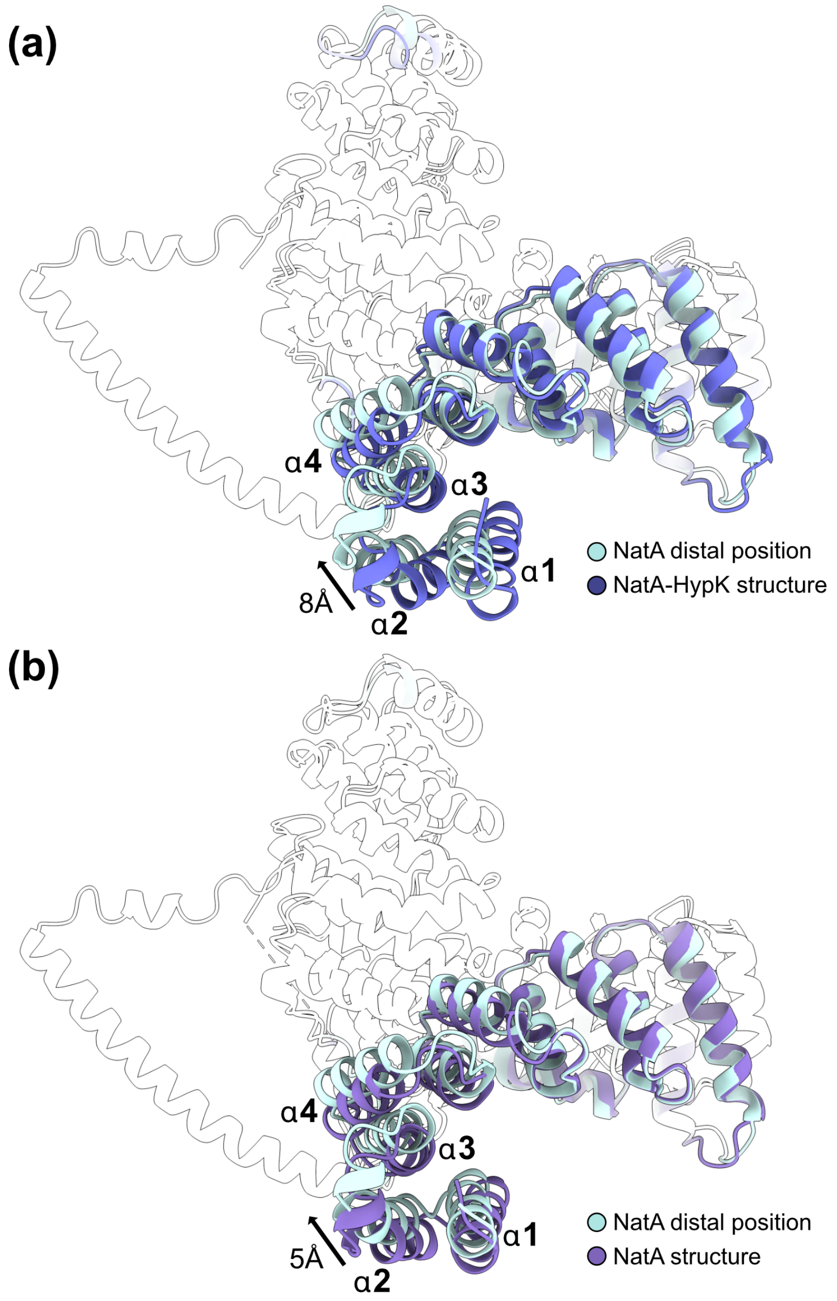


**Supplementary Figure 5: The helical Naa15 scaffold undergoes conformational changes to adopt the distal position on the ribosome. (a)** Compared to the structure of NatA-HypK[2] the N-termina helices shift by up to 8 Å. **(c)** Compared to the structure of NatA[2], this Naa15 rearrangement is also present, but less pronounced with a rotation of up to 5 Å. Naa15 binding of HypK or Naa15 binding to the distal position causes structural rearrangements of the N-terminal helices into opposing directions.

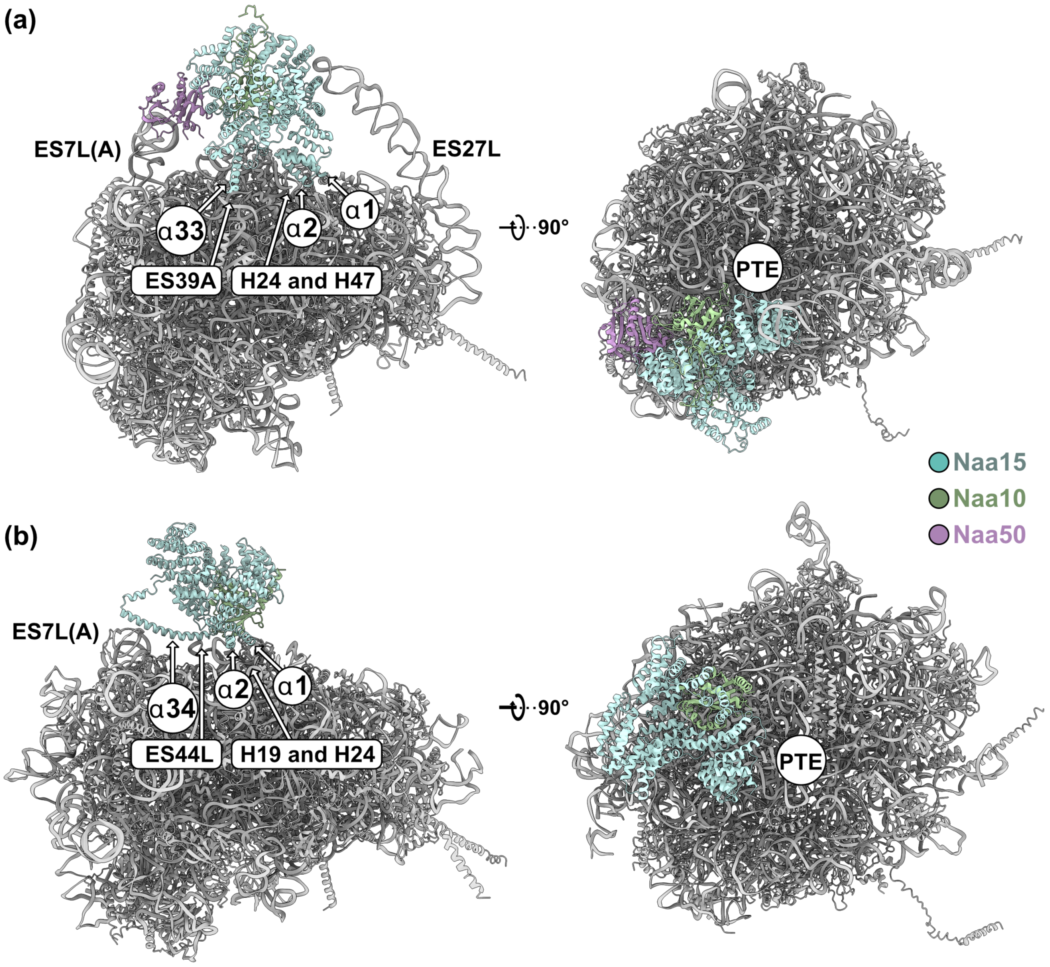


**Supplementary Figure 6: Comparison of the ribosomal binding site of *Hs*NatA and *Sc*NatE. (a)** Yeast NatE (PDB: 6HD7)[3] has four contact points with the 60S subunit of the ribosome. With contacts formed by helix α2, Naa15 is placed onto H24 and H47 directly adjacent to the PTE opening. The long anchoring helix α33 makes a second contact to ES39A. Mediated by Naa50, the yeast NatE complex contacts ES7L(A). The last contact is formed by the apex if ES27L and Naa15. **(b)** Binding of human NatA to the ribosome relies only on two contact points. The first contact is again mediated by the first two N-terminal helices α1 and α2 which touch down on H19 and H24. The second contact is mediated by the long anchoring helix α34 (corresponds to α33 in yeast) which wedges in between ES7L(A) and spans across the ribosomal surface to also interact with ES44L. The human NatA complex is positioned further away from the PTE (right panel).

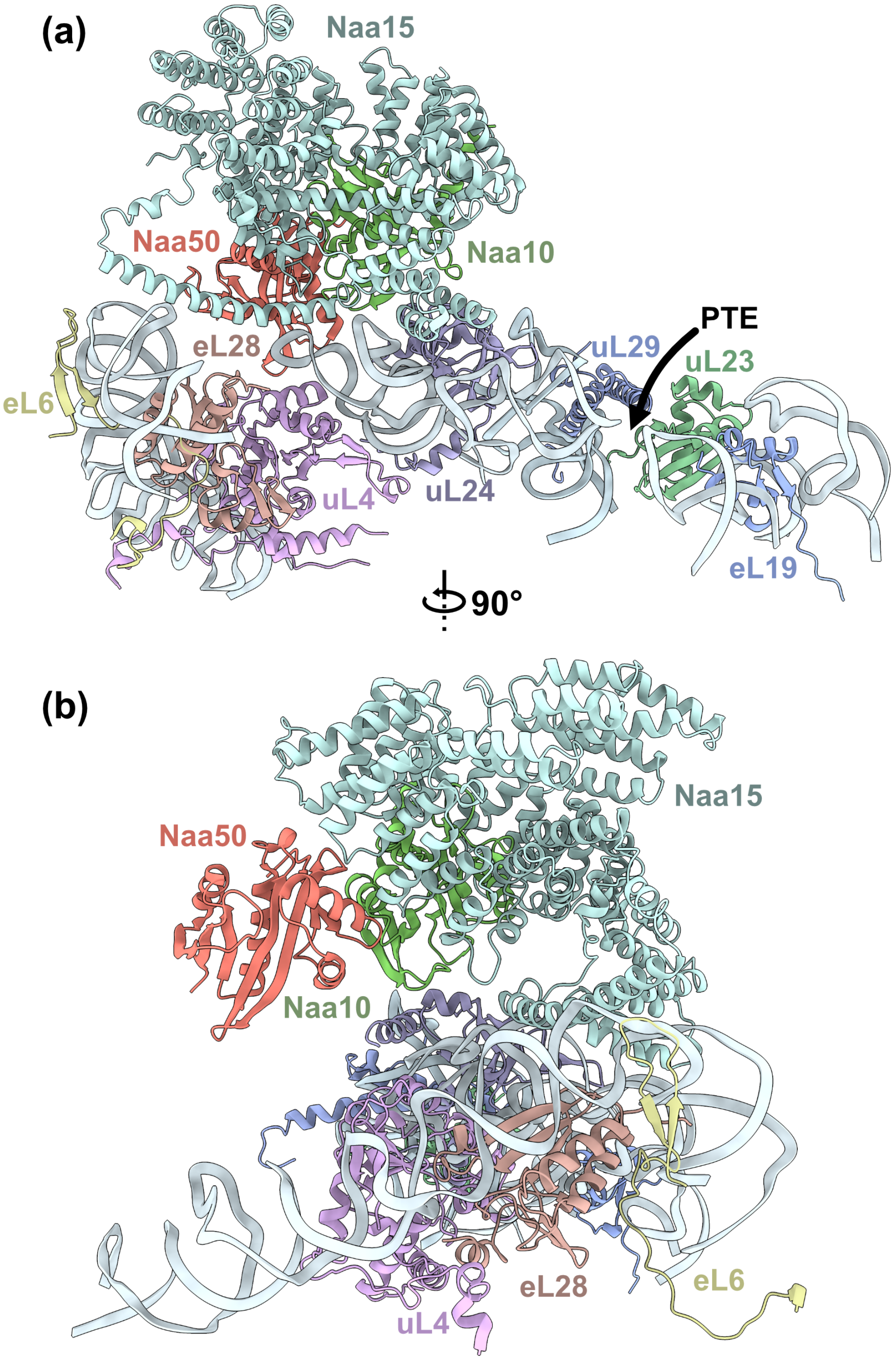


**Supplementary Figure 7: Crystal structure of human NatE superimposed on NatA in the distal position. (a)** Side view on NatA in the distal site. MAP2 is not shown and the position of the PTE is indicated by an arrow. Naa50 (red) would be positioned behind Naa15 and face away from the PTE (PDB: 6PPL was superimposed onto Naa15 in the distal site)[4]. **(b)** Top view onto NatA with Naa50 superimposed. In this position, Naa50 would not contact the ribosomal surface.

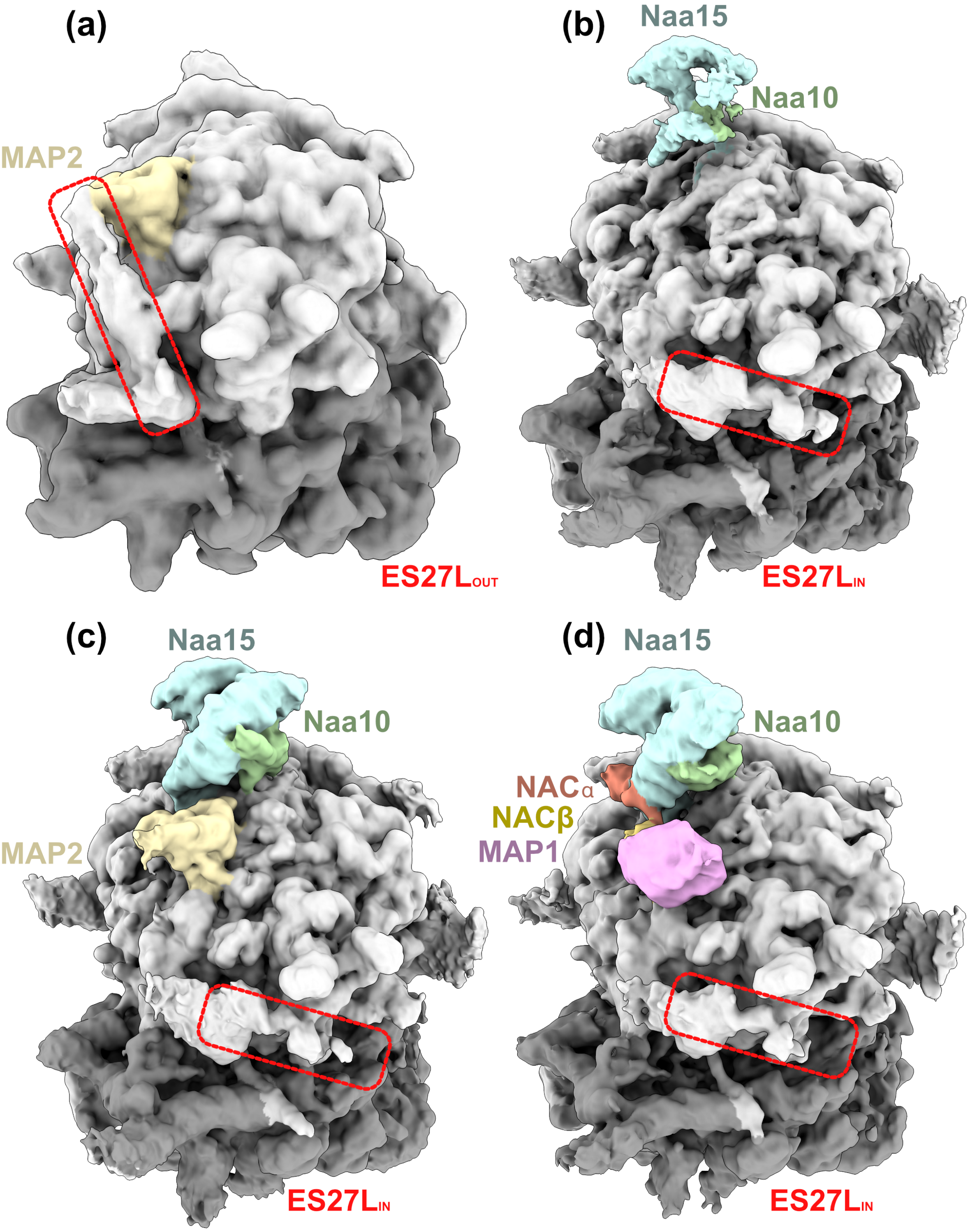


**Supplementary Figure 8: ES27L is not recruited to the ES27L$_{out}$ position on NatA decorated ribosomes. (a)** In complex with MAP2, ES27L is recruited to the ES27L$_{OUT}$ position on a subset of particles[5]. In complex with **(b)** NatA, **(c)** NatA-MAP2 and **(d)** NatA-NAC-MAP1, ES27L is placed in the ES27L$_{in}$ position.

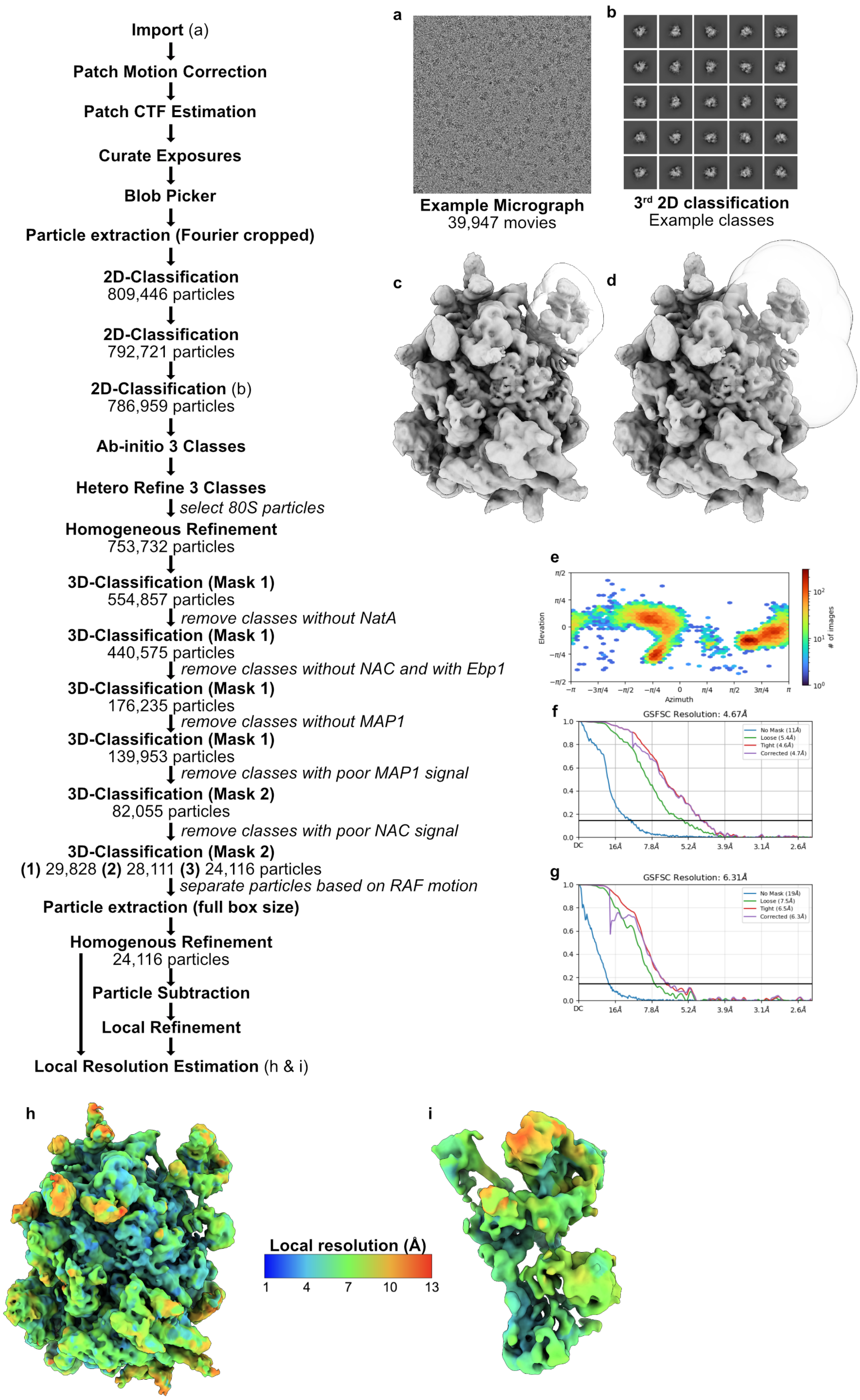


**Figure description is located on the next page**

**Supplementary Figure 9: Cryo-EM data processing for the *Hs*NatA-*Hs*NAC-*Hs*MAP1-80S sample. (a)** Three datasets were acquired to obtain 39,947 movies. **(b)** After pre-processing in CryoSPARC, extracted particles were subjected to three rounds of 2D-classification. Three ab-initio classes were generated from the remaining particles and used to seed a Heterogeneous refinement. 80S particles were selected and subjected to Homogenous Refinement. **(c)** A mask (Mask 1) was generated to encompass the distal site of NatA and used to initialize a 3D classification to remove particles without NatA. The same mask was used for three additional subsequent 3D classifications to remove particles with Ebp1, without NAC, without MAP1 or with poor MAP1 signal. **(d)** A second extended mask (Mask 2) was made to encompass the binding site of NatA, NAC and MAP1 and used to initialize a fourth 3D classification to remove particles with poor NAC signal. The same mask was finally used in a fifth 3D classification into three classes to separate particles based on the motion of NatA, NAC and MAP1. One of the classes revealed the complex in a more stabilized state. Corresponding particles were re-extracted at full box size without Fourier cropping and subjected to Homogenous refinement. **(e)** Angular distribution plot of the final Homogenous refinement. **(f)** FSC plots of the final Homogenous refinement. **(g)** Finally, as mask was generated to subtract the 80S signal and the mask shown in (d) was used to initialize a local refinement around MAP1, NAC and NatA. The corresponding FSC curve is shown. **(h)** Local resolution estimation of the final Homogenous refinement. **(i)** Local resolution estimation of the final local refinement.

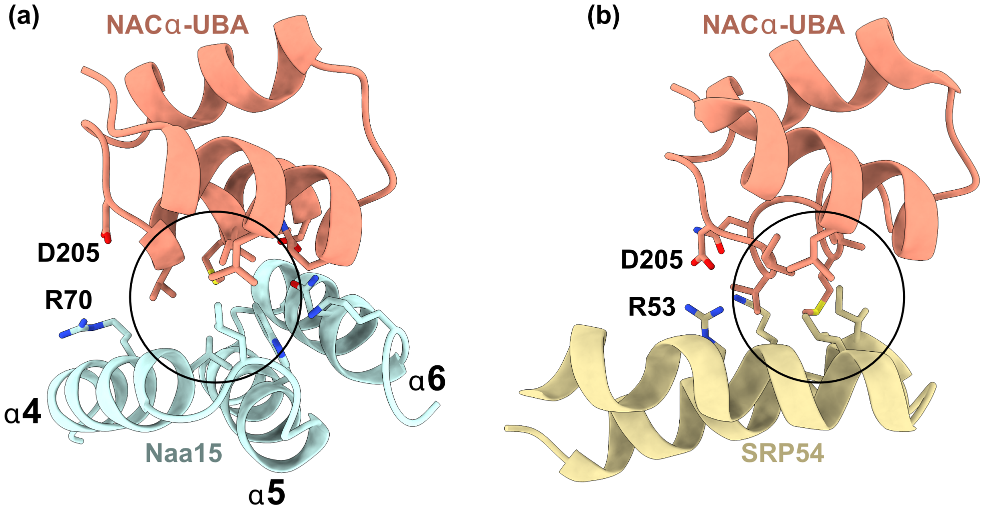


**Supplementary Figure 10: The NACα UBA domain can contact Naa15 and SRP54. (a)** The NAC UBA domain can contact helices α4-6 with an exposed hydrophobic patch. Surrounding polar and ionic interactions further stabilize the complex. **(b)** The interaction between the NACα-UBA domain and SRP54 is also mediated by a hydrophobic core (Figure made from PDB entry 7QWQ)[6]. D205$^{NAC\alpha}$ appears to form an ionic interaction on the surface of Naa15 and SRP54.

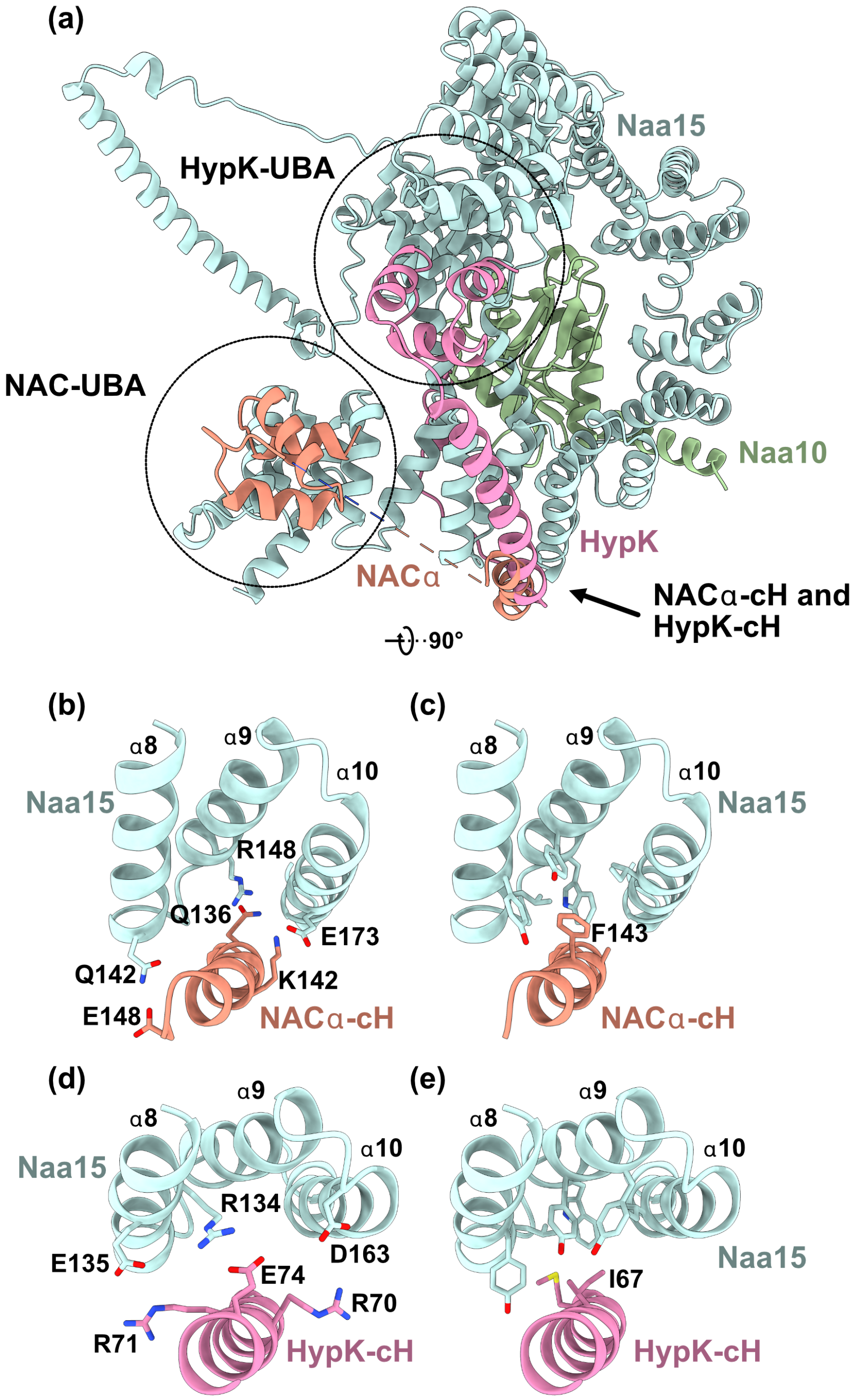


**Supplementary Figure 11: Homologues HypK and NACα form similar interactions with Naa15. (a)** Interaction between the C-terminus of NACα (including NACα-cH and UBA domain) superimposed with HypK (from pdb entry 6C95[2]). NACα-cH and HypK-cH both run along the surface of Naa15 helices α8-10. The HypK-cH helix is longer than the NACα-cH. While the NACα-UBA domain is placed on top of helices α4-6 near the N-terminus of Naa15, the HypK-UBA domain is positioned near the C-terminal helices of Naa15. **(b)** The NACα-cH runs along the surface of Naa15 in parallel to helices α8-10 and engages in several polar and ionic interactions around a **(c)** central hydrophobic core. **(d)** The interaction of HypK and Naa15 around helices α8-10 of Naa15 is mainly stabilized by ionic interactions that **(e)** surround the central hydrophobic core.

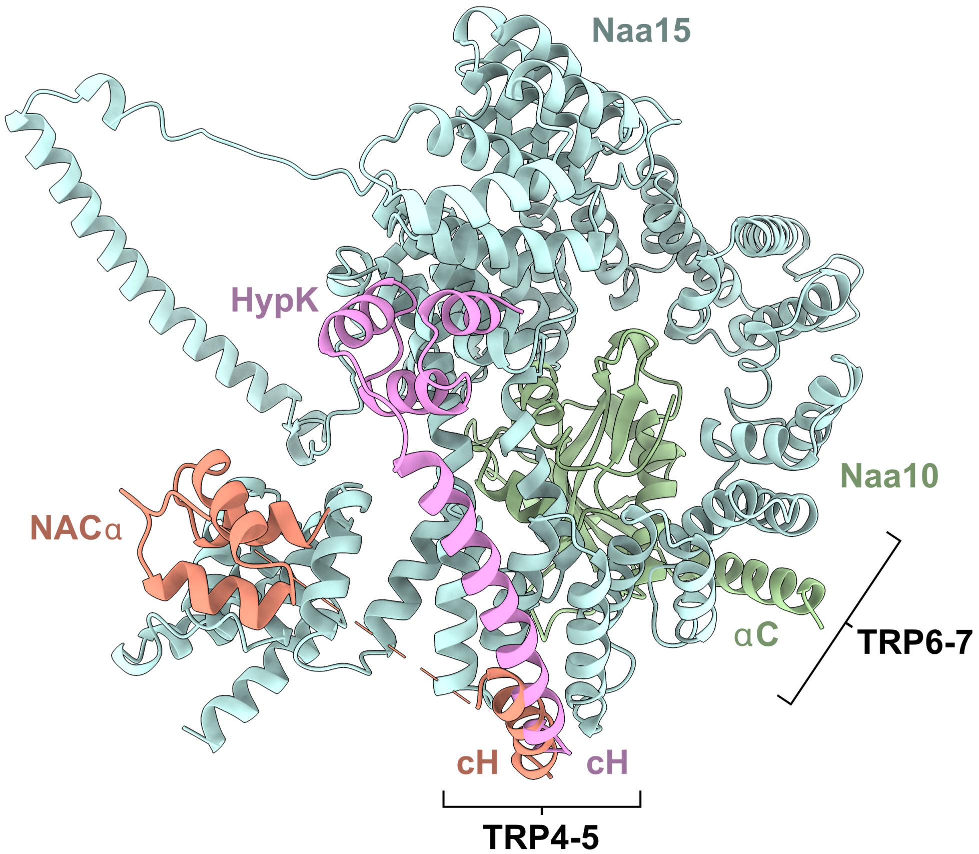


**Supplementary Figure 12: The Naa15 scaffold engages in helical contacts with Naa10-αC, NACα-cH and HypK-cH.** The C-terminal helix of Naa10 (αC) runs along the surface of Naa15 TPR6 and TPR7 (helices α12-14). This parallel placement of the Naa10 C-terminal helix resembles the interaction of the NACα-cH and HypK-cH helix with TPR4 and TPR5 of Naa15 (helices α8-10).

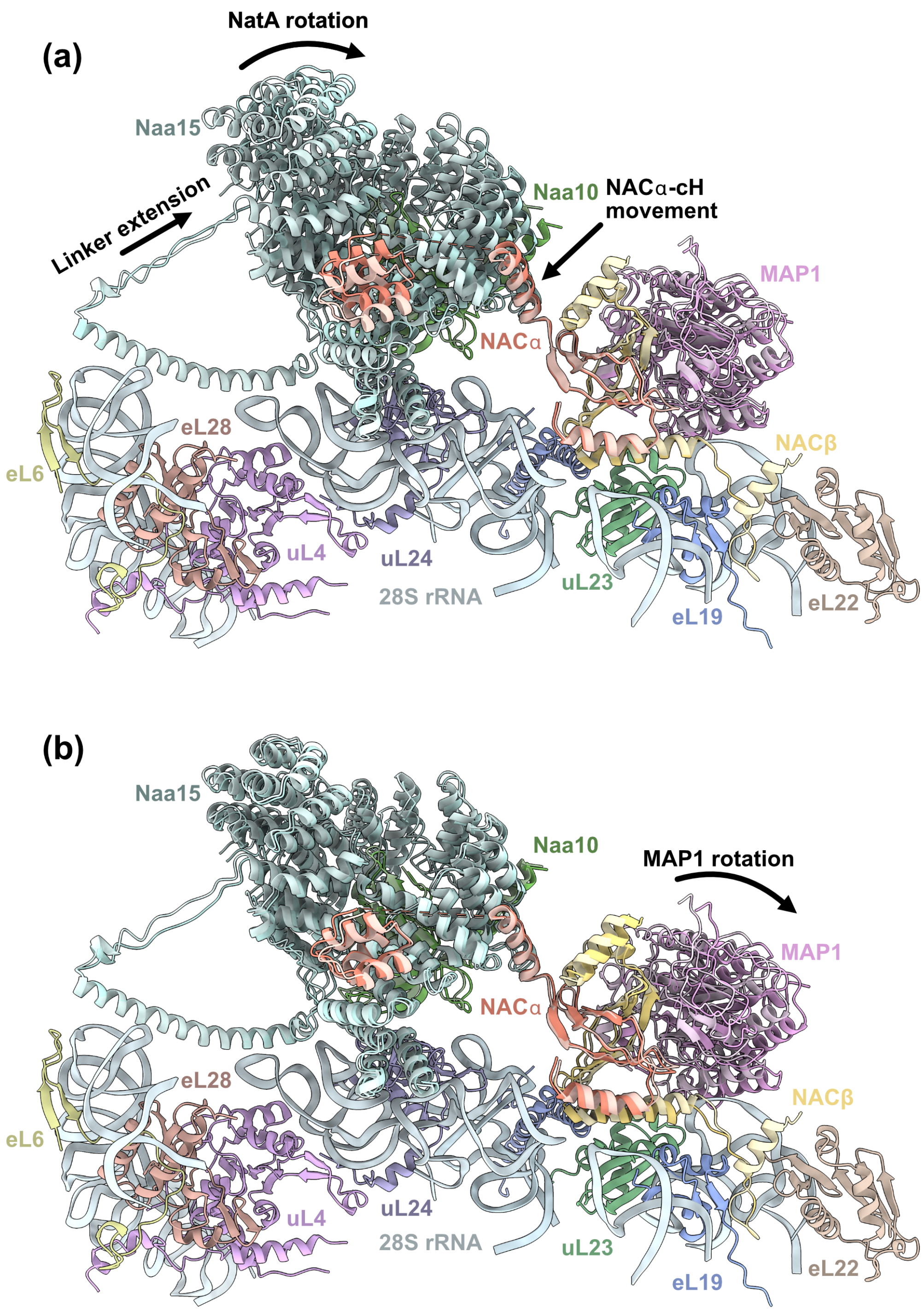


**Supplementary Figure 13: Variability of the quaternary NatA-NAC-MAP1-80S assembly.** In the final 3D classification, particles were separated into three classes (Classes 1-3) with different orientations of NatA, NAC and MAP1. **(a)** Comparison of Class 1 (desaturated colour) with Class 2 (saturated colour). In Class 2, NatA is rotated further down towards the PTE. The linker that connects NatA to the $\alpha$34 anchor is extended and the NAC$\alpha$-cH and UBA domain are shifted. **(b)** Comparison of Class 1 (desaturated colour) with Class 3 (saturated colour). In Class 3, the interaction between NAC and MAP1 is weaker and MAP1 is located further away from the PTE. The NAC dimerization domain is also slightly shifted.

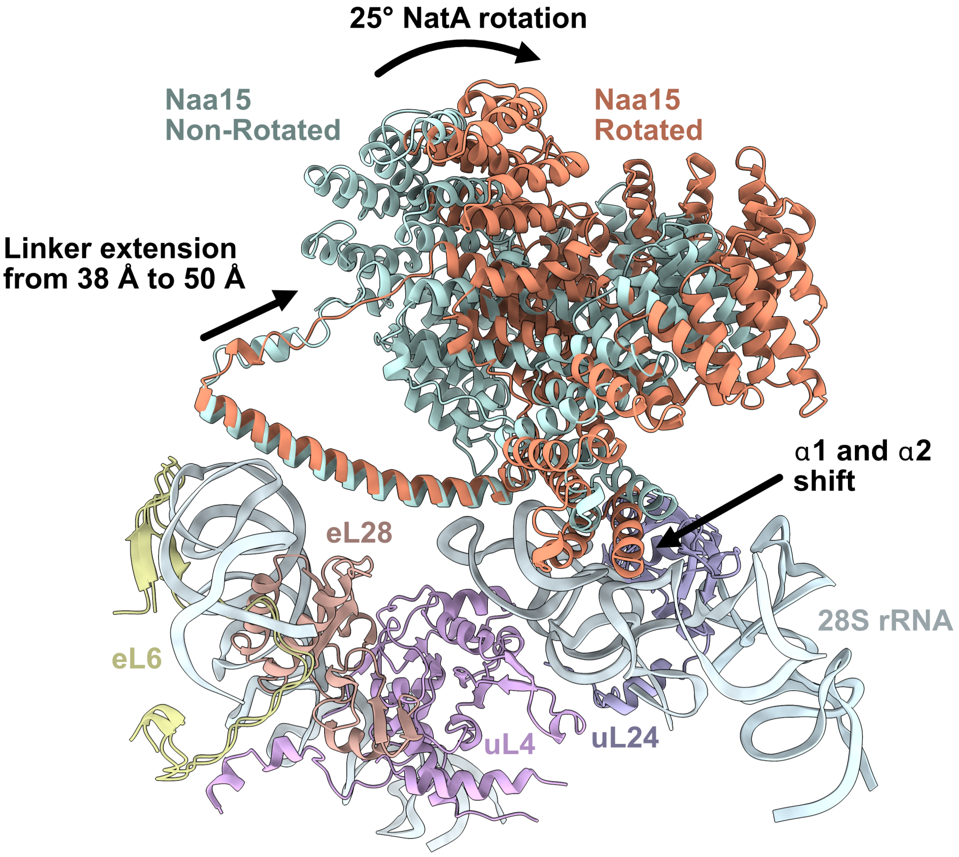


**Supplementary Figure 14: Differences in NatA binding in the ternary NatA-MAP2-80S complex compared to the quaternary NatA-NAC-MAP1-80S complex.** In complex with NAC and MAP1, the NatA complex is rotated further down towards the PTE (red). In this rotation Naa15 remains anchored in place. The contacts of Naa15 helices $\alpha$1 and $\alpha$2 are shifted further away from uL24 in the quaternary assembly. Throughout the rotation, a flexible linker can extend between 38 Å and 50 Å to enable this dynamic rotation of NatA by 25°.

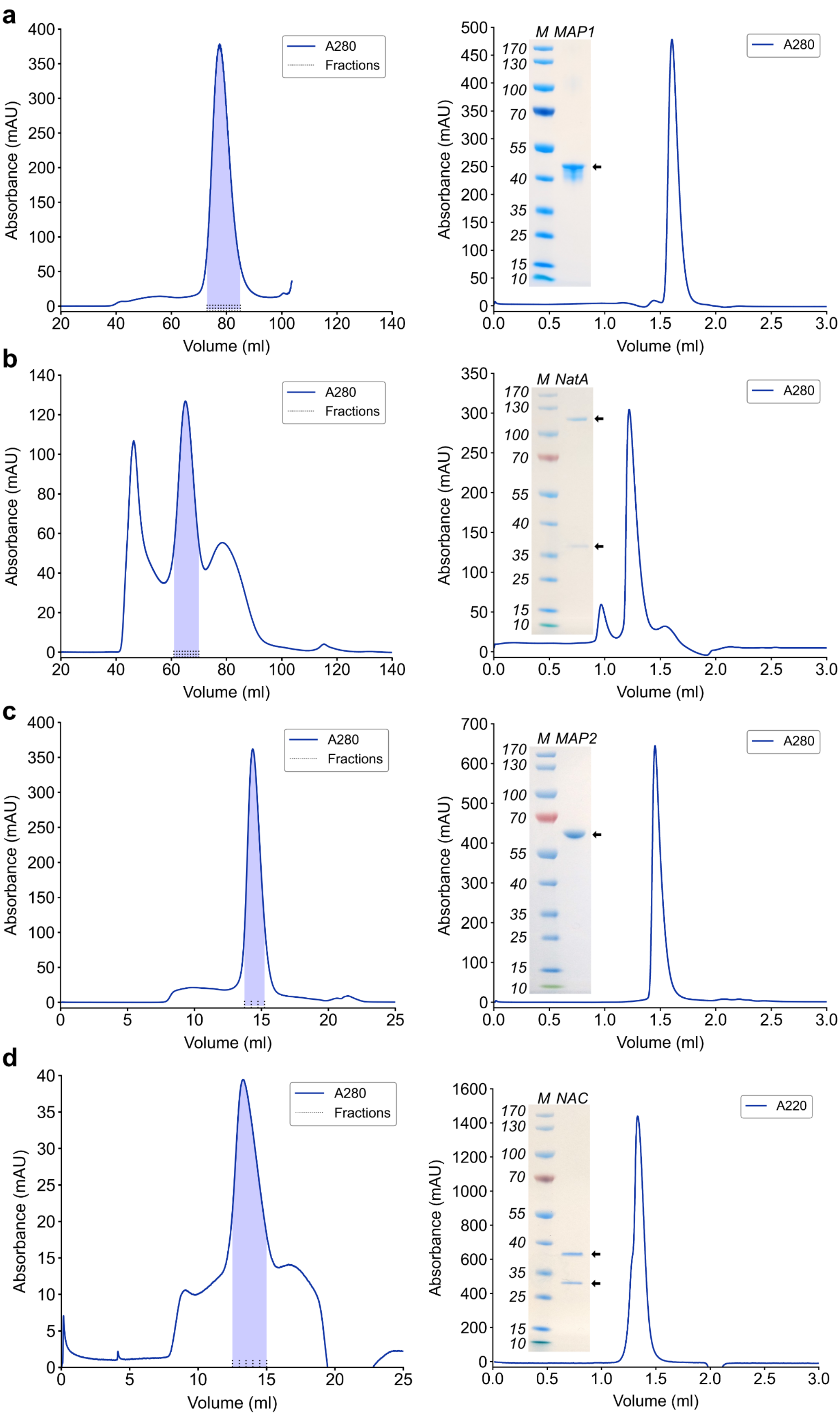


**Figure description is located on the next page**

**Supplementary Figure 15: Protein purifications of ribosome associated factors.** Pooled fractions from preparative SEC (shaded in blue) were subjected to SDS-PAGE and coomassie staining, as well as analytical SEC. Preparative SEC was performed on an S200 16/600 column (Cytiva) for **(a)** MAP1 and **(b)** NatA and on an S200 Increase 10/300 GL column (Cytiva) for **(c)** MAP2 and **(d)** NAC. Analytical SEC was done on an S200 Increase 3.2/300 column (Cytiva).

### **Supplementary Table 1:** Cryo-EM data refinement statistics

| **Model** | ***Hs*NatA-*Hs*MAP2-80S** | ***Hs*NatA-*Hs*NAC-*Hs*MAP1-80S** |
|---|---|---|
| *Data collection statistics* | | |
| Microscope | Titan Krios | Glacios |
| Camera | K3 | Falcon 3 |
| Voltage (kV) | 300 | 200 |
| Magnification | 105,000 | 120,000 |
| Total dose ($e^-/Å^2$) | 41.28 $e^-/Å^2$ | 53.50, 53.97 and 53.96 $e^-/Å^2$ |
| Defocus range ($\mu$m) | -1.3 to -2.3 | -0.7 to -1,7 |
| Calibrated pixel size (Å) | 0.84 | 1.223 |
| *Refinement statistics* | | |
| Refined particles | 25,404 | 24,116 |
| Resolution (Å) | 2.69 | 4.67 |
| Chains | 15 | 18 |
| Atoms | 29394 | 31688 (Hydrogens: 0) |
| Residues | Protein: 2666 Nucleotide: 358 | Protein: 2963 Nucleotide: 356 |
| Water | 0 | 0 |
| Ligands | IHP: 1<br>CO: 2 | IHP: 1 |
| Bonds (RMSD*) | | |
| Length (Å) (# > 4$\sigma$) | 0.003 (0) | 0.005 (0) |
| Angles (°) (# > 4 $\sigma$) | 0.740 (24) | 1.109 (65) |
| MolProbity score | 1.98 | 2.56 |
| Clash score | 14.75 | 30.73 |
| Ramachandran plot (%) | | |
| Outliers | 0.08 | 0.58 |
| Allowed | 3.14 | 4.30 |
| Favored | 96.78 | 95.12 |
| Rama-Z (Ramachandran Plot, Z-score, RMSD*) | | |
| whole (N = 951) | 0.42 (0.16) | -0.03 (0.15) |
| helix (N = 379) | 0.60 (0.14) | 0.37 (0.14) |
| sheet (N = 107) | -0.02 (0.38) | -0.49 (0.31) |
| loop (N = 465) | 0.12 (0.19) | -0.17 (0.18) |
| Rotamer outliers (%) | 1.36 | 2.11 |
| C$\beta$ outliers (%) | NA | 0.07 |
| Peptide plane (%) | | |
| Cis proline/general | 0.0/0.0 | 1.7/0.0 |
| Twisted proline/general | 0.0/0.0 | 0.8/0.0 |
| CaBLAM outliers (%) | 1.45 | 1.76 |
| ADP (B-factors) | | |
| Iso/Aniso (#) | 29394/0 | 31688/0 |
| min/max/mean | | |
| Protein | 14.93/495.28/134.23 | -0.00/990.92/204.79 |
| Nucleotide | 66.29/489.52/152.28 | 2.28/798.14/216.25 |
| Ligand | 115.67/250.44/122.56 | 235.80/235.80/235.80 |
| Occupancy (%) | | |
| Mean | 1.00 | 1.00 |
| occ = 1 (%) | 100.00 | 100.00 |
| 0 < occ < 1 (%) | 0.00 | 0.00 |
| occ > 1 (%) | 0.00 | 0.00 |
| Model vs. Data (CC mask) | 0.80 | 0.53 |
| Resolution according to model vs. map FSC = 0.143 (masked) (Å) | 2.9 | 4.8 |

* RMSD: root-mean-squared-deviation
